# Dystroglycan N-terminal domain enables LARGE1 to extend matriglycan on α-dystroglycan and prevents muscular dystrophy

**DOI:** 10.1101/2022.08.09.502851

**Authors:** Hidehiko Okuma, Jeffrey M. Hord, Ishita Chandel, David Venzke, Mary E. Anderson, Ameya S. Walimbe, Soumya Joseph, Zeita Gastel, Yuji Hara, Fumiaki Saito, Kiichiro Matsumura, Kevin P. Campbell

## Abstract

Dystroglycan (DG) requires extensive post-translational processing to function as a receptor for extracellular matrix proteins containing laminin-G-like (LG) domains. Matriglycan is an elongated polysaccharide of alternating xylose and glucuronic acid that is uniquely synthesized on α-dystroglycan (α-DG) by like-acetylglucosaminyltransferase-1 (LARGE1) and binds with high affinity to matrix proteins like laminin. Defects in the post-translational processing of α-DG that result in a shorter form of matriglycan reduce the size of α-DG and decrease laminin binding, leading to various forms of muscular dystrophy. However, little is known regarding mechanisms that generate full-length matriglycan on α-DG (~150-250 kDa). Here, we show that LARGE1 can only synthesize a short, non-elongated form of matriglycan in mouse skeletal muscle that lacks the DG N-terminus (α-DGN), resulting in a ~100-125 kDa α-DG. This smaller form of α-DG binds laminin and maintains specific force but does not prevent muscle pathophysiology, including reduced force induced by eccentric contractions and abnormalities in neuromuscular junctions. Collectively, our study demonstrates that α-DGN is required for LARGE1 to extend matriglycan to its full mature length on α-DG and thus prevent muscle pathophysiology.

## Introduction

The basement membrane is a specialized network of extracellular matrix macromolecules that surrounds epithelium, endothelium, muscle, fat, and neurons (***Rowe and Weiss, 2008***). Skeletal muscle cells are bound to the basement membrane through transmembrane receptors, including dystroglycan (DG) and the integrins, which help maintain the structural and functional integrity of the muscle cell membrane (***Roberts et al., 1985; Sonnenberg et al., 1988; Ibraghimov-Beskrovnaya et al., 1992; Han et al., 2009***). DG is a central component of the dystrophin-glycoprotein complex (DGC). It is encoded by a single gene, DAG1, and cleaved into α- and β-subunits (α-DG and β-DG, respectively) by post-translational processing (***Ibraghimov-Beskrovnaya et al., 1992***). Extensive *O*-glycosylation of α-dystroglycan (α-DG) is required for normal muscle function, and defects in this process result in various forms of muscular dystrophy. (***Michele et al., 2002; Yoshida-Moriguchi and Campbell, 2015***)

α-DG binds to ECM ligands containing laminin-G domains (e.g., laminin, agrin, perlecan) that are essential components of the basement membrane (***Michele et al., 2002***). DG, therefore, physically links the cell membrane to the basement membrane. This process requires synthesis of matriglycan, a heteropolysaccharide [-GlcA-β1,3-Xyl-α1,3-]_n_, on α-DG by the bifunctional glycosyltransferase, like-acetylglucosaminyltransferase-1 (LARGE1) (***Chiba et al., 1997; Michele et al., 2002; Inamori et al., 2012; Yoshida-Moriguchi and Campbell, 2015; Hohenester, 2019; Michele et al., 2002; Ohtsubo and Marth, 2006***). *O*-glycosylation and the glycosylation-specific kinase, Protein O-Mannose Kinase (POMK), which phosphorylates mannose of the core M3 trisaccharide (GalNAc-β1,3-GlcNAc-β1,4-Man), are required to produce full-length, high-molecular weight forms of matriglycan (***Yoshida-Moriguchi and Campbell, 2015; Hohenester, 2019; Jae et al., 2013; Yoshida-Moriguchi et al., 2013; Zhu et al., 2016***). In the absence of phosphorylation of core M3 by POMK, LARGE1 synthesizes a short, non-elongated form of matriglycan on α-DG (***Walimbe et al., 2020***). Notably, a loss of function in the post-translational addition of matriglycan causes dystroglycanopathies, which are congenital and limb-girdle muscular dystrophies with or without brain and eye abnormalities. Disease severity is dependent on the ability of matriglycan to bind ECM ligands, which is dictated by its length and expression (***Goddeeris et al., 2013***): matriglycan that is low molecular weight (e.g., short) can cause muscular dystrophy, even if its capacity to bind laminin-G domains is not completely lost (***Puckett et al., 2009; Hara et al., 2011; Carss et al., 2013; Cirak et al., 2013; Dong et al., 2015; Walimbe et al., 2020***). However, the regulation of matriglycan elongation by factors other than POMK is still unknown.

α-DG is composed of three distinct domains: the N-terminal (α-DGN) domain, a central mucin-like domain, and the C-terminal domain (***Figure 1***) (***Brancaccio et al., 1995***). α-DGN functions as a binding site for LARGE1 in the Golgi and is required for the functional glycosylation of the mucin-like domain of α-DG (***Kanagawa et al., 2004***). Therefore, we hypothesized that α-DGN must be involved in regulating the production and elongation of matriglycan. Here, we used a multidisciplinary approach to show that LARGE1 synthesizes a non-elongated form of matriglycan on DG that lacks α-DGN (i.e., α-DGN-deleted dystroglycan) resulting in ~100-125 kDa α-DG. This short form of matriglycan binds laminin and maintains muscle-specific force. However, it fails to prevent lengthening contraction-induced reduction in force, neuromuscular junction abnormalities, or dystrophic changes in muscle, as these effects require the expression of α-DG with the matriglycan modification that is at least 150 kDa. Therefore, this study shows that LARGE1 requires α-DGN to generate full-length (~150-250 kDa) matriglycan in skeletal muscle, but synthesis of a shorter form of matriglycan still occurs in the absence of this domain.

**Figure 1.**
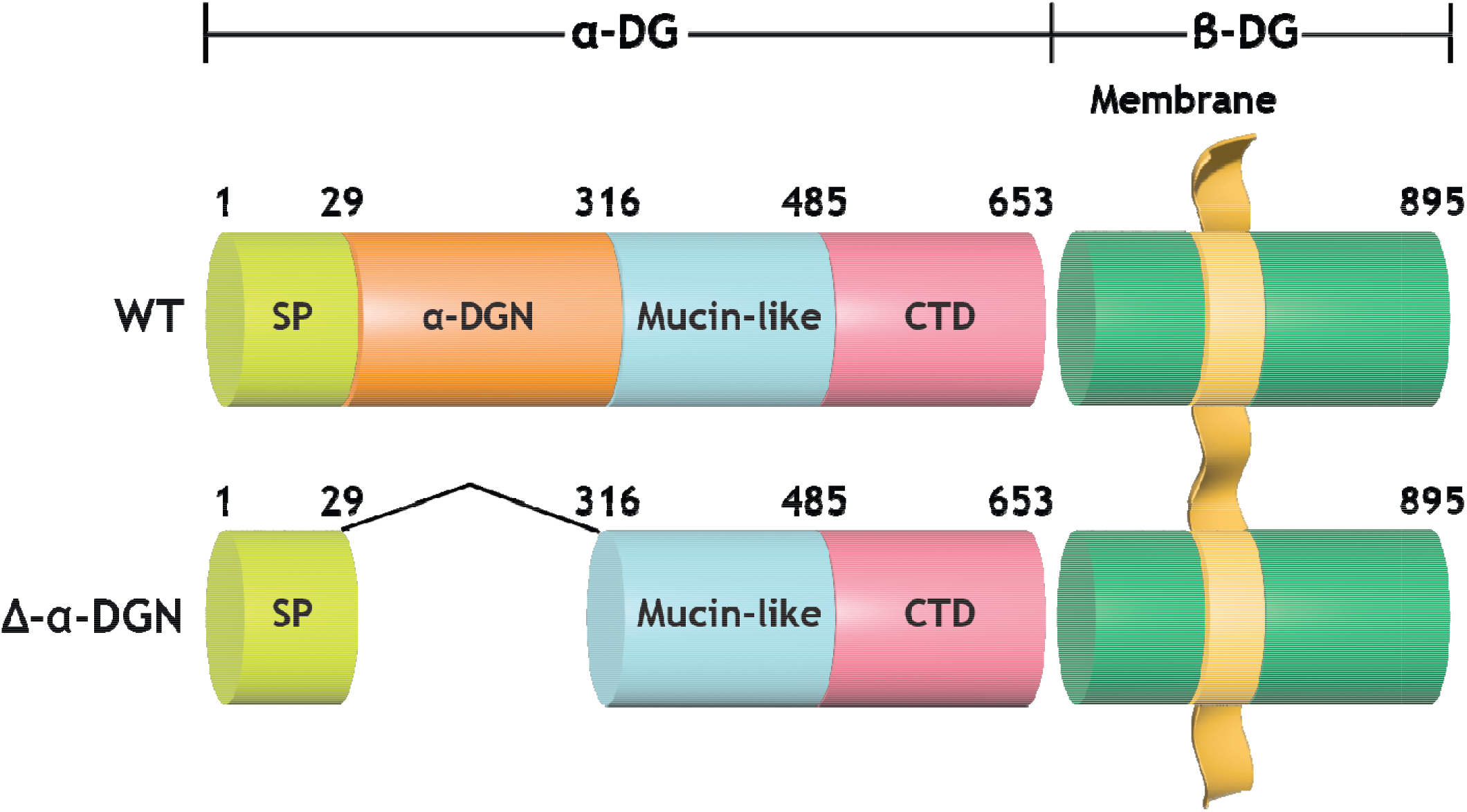
Domain structure of DG and A-α-DGN. Wild-type DG is a pre-proprotein with an N-terminal signal peptide (light green) that is translated in the rough endoplasmic reticulum. The globular N-terminal domain (α-DGN; orange) is present in wild-type DG but absent in the mutant (Δ-α-DGN). The junction between α-DGN and the mucin-like domain (light teal) contains a furin convertase site. The globular extracellular C-terminal domain (CTD; pink) contains a SEA (sea urchin sperm protein, enterokinase and agrin) autoproteolysis site, which cleaves pro-DG into α-DG and β-DG (green). Glycosylation has been omitted for clarity.

## Results

To ablate α-DGN in skeletal muscle, we used mice expressing Cre recombinase under the control of the *paired box 7* (*Pax7*) promoter (*Pax7*^Cre^), floxed DG mice (*Dag1^flox/flox^*), and heterozygous α-DGN deleted mice (*Dag1^wt/Δα-DGN^*) to generate *Pax7^Cre^Dag1^flox/Δα-DGN^* (M-α-DGN KO) mice (***Figure 2***). Constitutive deletion of DG in mice causes embryonic lethality due to the absence of Reichert’s membrane, an extraembryonic basement membrane required for *in utero* development (***Williamson et al., 1997***). Deletion of α-DGN in mice also causes embryonic lethality (***de Greef et al., 2019***). However, mice that are heterozygous for α-DGN deletion are viable and express α-DG of two different sizes (***Figure 2-figure supplement 1***) corresponding to both wild-type (WT) and the α-DGN-deleted (Δα-DGN) forms of DG.

**Figure 2.**
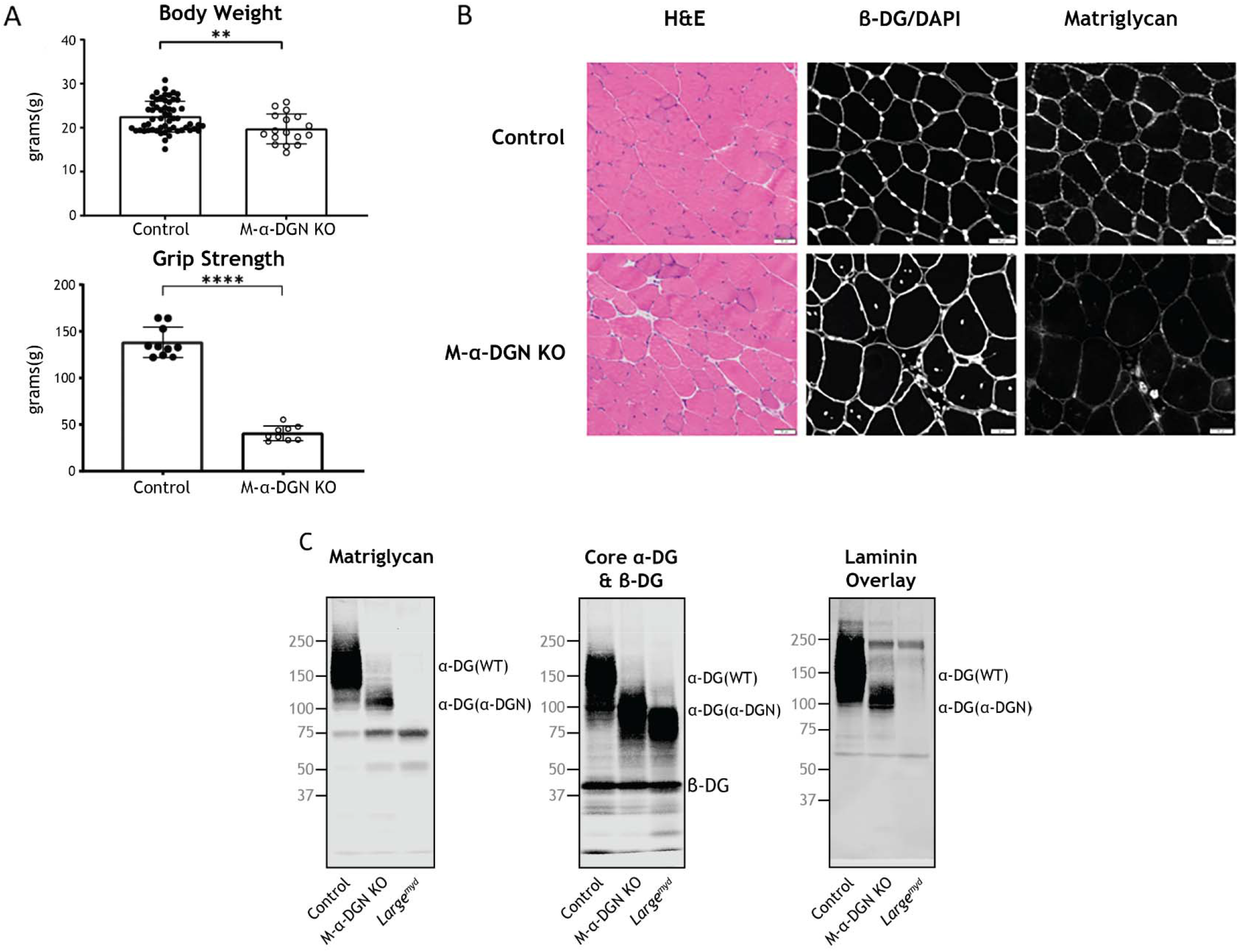
Characterization of muscle-specific α-DGN-deficient mice. **(A)** Body weight and grip strength of 12-week-old WT littermates (control) and M-α-DGN KO mice. Double and quadruple asterisks: statistical significance determined by Student’s unpaired t-test (**p-value=0.005, ****p-value<0.0001). **(B)** Histological analyses of quadriceps muscles from 12-week-old control and M-α-DGN KO mice. Sections stained with H&E or used for immunofluorescence to detect β-DG (affinity purified rabbit anti-β-DG), DAPI, and matriglycan (IIH6). **(C)** Immunoblot analysis of quadriceps skeletal muscle from control, M-α-DGN KO, and *Large^myd^* mice. Glycoproteins were enriched using wheat-germ agglutinin (WGA)-agarose with 10 mM EDTA. Immunoblotting was performed to detect matriglycan (IIIH11), core α-DG, β-DG (AF6868), and laminin overlay. α-DG in WT control muscle (α-DG (WT)) and α-DG in α-DGN-deficient muscle (α-DG (Δα-DGN)) are indicated on the right. Molecular weight standards in kilodaltons (kDa) are shown on the left.

To evaluate the gross phenotype of mice expressing only α-DGN-deleted DG in skeletal muscle (i.e., M-α-DGN KO mice), we first measured body weight and grip strength. M-α-DGN KO mice were lower in weight than WT littermates (control) mice at 12 weeks and they exhibited decreased forelimb grip strength (***Figure 2A***). To determine whether deletion of α-DGN affects matriglycan expression, we performed histological analysis of quadriceps muscle from control or M-α-DGN KO mice. M-α-DGN KO mice showed characteristic features of muscular dystrophy, including an increase in centrally nucleated fibers (***Figure 2B***). Immunofluorescence analyses of M-α-DGN KO muscle showed reduced levels of matriglycan relative to controls, but a similar expression of β-DG, the transmembrane subunit of DG (***Figure 2B***).

Immunoblot analysis of skeletal muscle from M-α-DGN KO mice demonstrated expression of a shorter form of matriglycan resulting in a ~100-125 kDa α-DG, a decrease in the molecular weight of the core α-DG, and no change in β-DG (***Figure 2C***). No matriglycan is seen in *Large^myd^* mice which have a deletion in *Large1 (**Figure 2C***). To investigate how the loss of α-DGN affected ligand binding, we performed a laminin overlay using laminin-111. Skeletal muscle from control mice showed a broad band centered at ~100-250 kDa, indicative of α-DG-laminin-binding; in contrast, we observed laminin-binding at ~100-125 kDa in M-α-DGN KO skeletal muscle (***Figure 2c***). To further confirm that the ~100-125 kDa band seen with anti-matriglycan antibodies in M-α-DGN KO muscle is matriglycan, we digested it overnight with β-glucuronidase (*Thermotoga maritima*) and α-xylosidase (*Sulfolobus solfataricus*). Immunoblot analysis after digestion with anti-matriglycan antibodies or laminin overlay revealed that the ~100-125 kDa was completely lost, indicating that the ~100-125 kDa band is indeed matriglycan on α-DG (***Figure 2-figure supplement 2***).

The neuromuscular junctions (NMJs) in adult control mice showed a normal pretzel-like shape whereas NMJs from M-α-DGN KO mice displayed a variety of abnormalities, including a granular appearance and AChR-rich streaks extending beyond the pre-synaptic terminal (***Figure 3***). Postsynaptic morphology in adult M-α-DGN KO mice was predominately irregular in the tibialis anterior (TA), extensor digitorum longus (EDL), and soleus (SOL) muscles (***Figure 3***). Although the overall synaptic size did not differ between controls and M-α-DGN KO mice, the dispersion of AChR clusters was greater in the M-α-DGN KOs (***Figure 3***), in line with an increased percentage of plaque-like formations and AChR extensions that projected beyond the nerve terminal. Despite the post-synaptic abnormalities, all NMJs from M-α-DGN KO mice were fully innervated.

**Figure 3.**
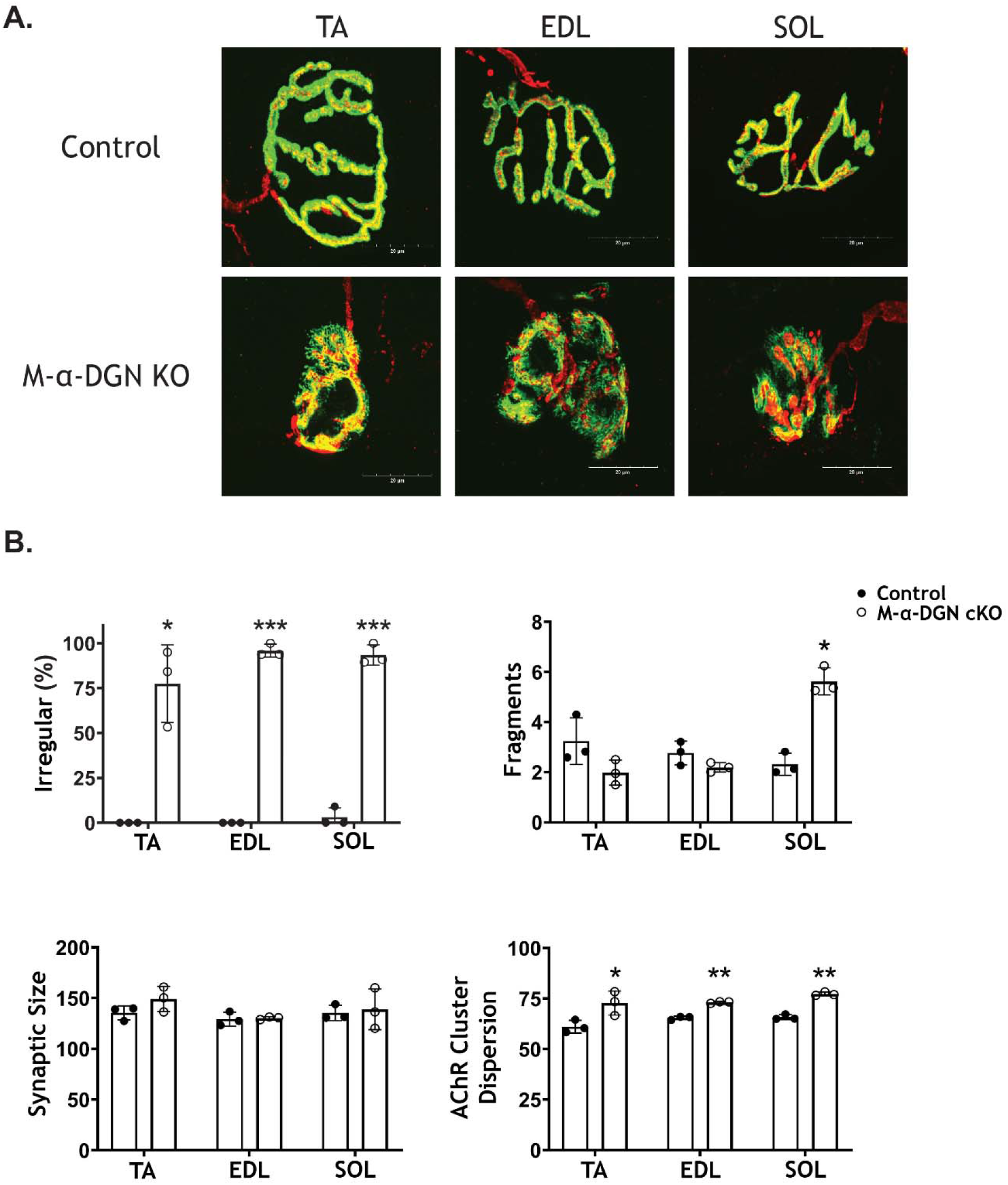
α-DGN deficiency results in postsynaptic defects. Neuromuscular junctions (NMJs) from tibialis anterior (TA), extensor digitorum longus (EDL), and soleus (SOL) muscles obtained from 35-39-week-old adult control and M-α-DGN KO mice. **(A)** Representative images of post-synaptic terminals (α-BTX-488; green), motor axons (anti-neurofilament-H; red), and pre-synaptic terminals (anti-synaptophysin; red) from TA, EDL, and SOL muscles. Scale bars = 20 μm. **(B)** Scoring of postsynaptic defects by blinded observers (scoring criteria described in Methods). Statistical significance determined by Student’s unpaired t-test; * p-value < 0.05; ** p-value < 0.001; *** p-value < 0.0001.

To determine the effect that the loss of α-DGN has on muscle force production, we characterized the phenotype and function of EDL muscles in 12-17-week-old WT (control) and M-α-DGN KO mice. Specifically, we measured muscle mass, muscle cross-sectional area (CSA), production of absolute isometric tetanic force, specific force, and lengthening contraction-induced reduction in force. Muscle mass and CSA were comparable between control and M-α-DGN KO mice (***Figure 4A and B***). Although the production of absolute isometric tetanic force was significantly lower in M-α-DGN KO mice than in control mice (***Figure 4C***), specific forces were comparable between the two groups when normalized to muscle CSA (***Figure 4D***). Lengthening contraction-induced reduction in force for M-α-DGN KO EDL remained greater than those from control EDL for the entire 60 minutes that muscles were assessed (***Figure 4E***). These results suggest that the short form of matriglycan on α-DG in M-α-DGN KO mice enables force production but cannot prevent force reduction caused by lengthening contractions. POMK KO skeletal muscle also expresses a short form of matriglycan, similar to M-α-DGN KO muscle, which maintains force production but cannot prevent lengthening contraction-induced force decline (***Walimbe et al., 2020***). Therefore, we compared the muscle function of EDL muscles from POMK KO mice with those from M-α-DGN KO *ex vivo*. We did not observe significant differences in lengthening contraction-induced force deficits between the two mouse strains (***Figure 4-figure supplement 1***). These results suggest that matriglycan of a similar molecular weight exhibits similar muscle force production and lengthening contraction-induced force decline.

**Figure 4.**
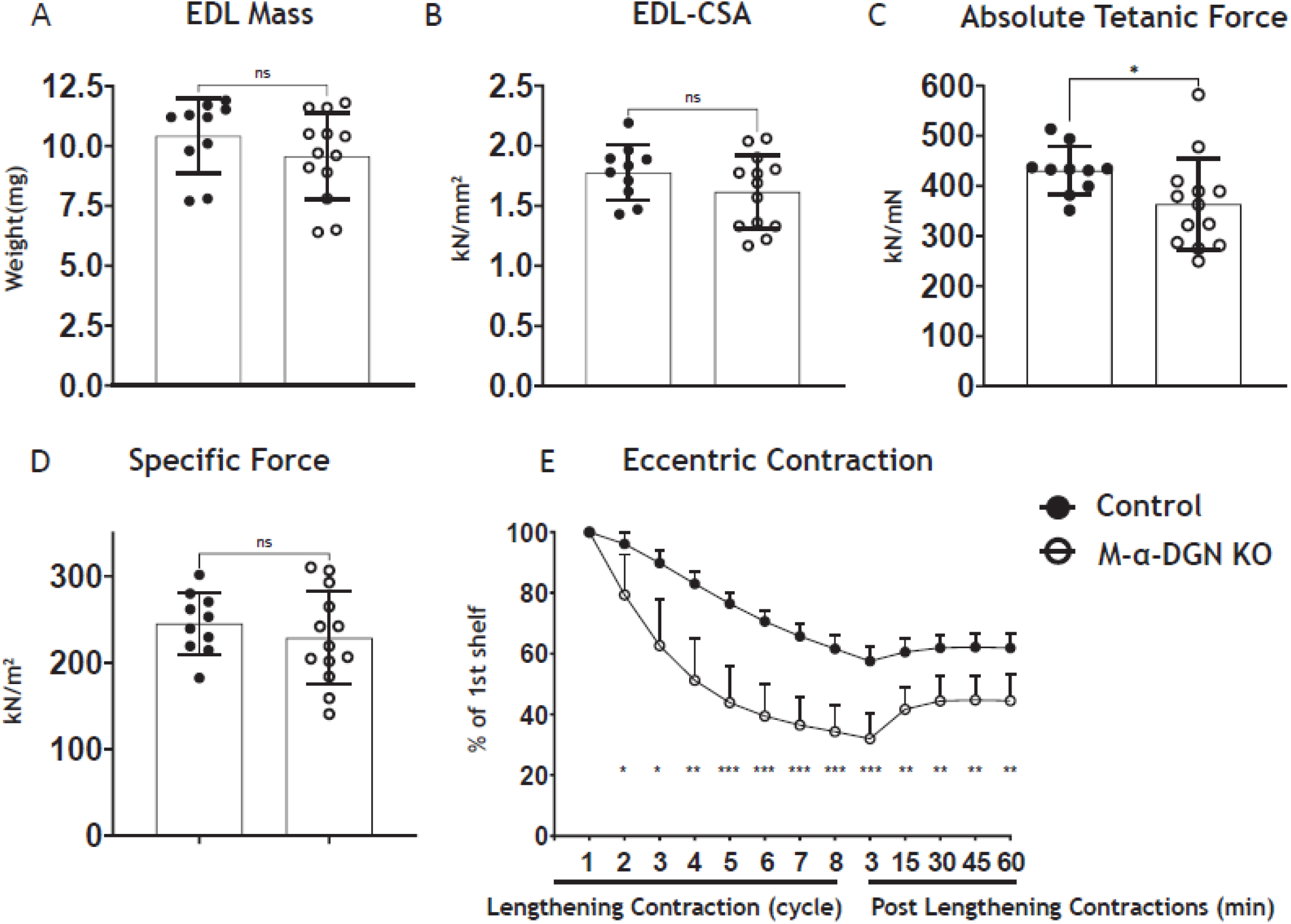
α-DGN-deficient Extensor Digitorum Longus (EDL) muscle demonstrates a decline in lengthening contraction-induced force. **(A)** Weight (milligrams) of EDL muscles from WT littermates (controls) and M-α-DGN KO mice; p=0.2469, as determined by Student’s unpaired t-test. **(B)** Cross-sectional area of EDL muscles; p=0.1810, as determined by Student’s unpaired t-test. **(C)** Maximum absolute tetanic force production in EDL muscles. p=0.0488, as determined by Student’s unpaired t-test. **(D)** Specific Force production in EDL muscles; p=0.4158, as determined by Student’s unpaired t-test. **(E)** Force deficit and force recovery after eccentric contractions in EDL muscles from 12- to 17-week-old male & female control (closed circles; n=7) and M-α-DGN KO (open circles; n=7) mice. *p<0.05; **p<0.01; ***p<0.001, as determined by Student’s unpaired t-test of at any given lengthening contractions cycle. Bars represent the mean +/- the standard deviation.

We next determined if exogenous DG lacking α-DGN (***Figure 5***) produces the short form of matriglycan. We first produced muscle-specific DG KO mice to achieve muscle-specific deletion of DG. To do this, we used mice expressing Cre under control of the paired box 7 (Pax7) promoter (Pax7-Cre) and *Dag1^flox/flox^* mice to generate Pax7Cre; *Dag1^flox/flox^* (M-*Dag1* KO) mice. To assess muscle function, we evaluated muscle-specific force and lengthening contraction-induced reduction in force *ex vivo*, which showed that muscle-specific force was significantly decreased and that muscles were more susceptible to lengthening contraction-induced force decline in the absence of DG (***Figure 5-figure supplement 1***). Collectively, these results show that M-*Dag1* KO mice harbor a more complete deletion of DG in muscle than the previously generated mouse model (MCK-Cre *Dag1^flox/flox^*) harboring muscle-specific deletion of DG (***Cohn et al., 2002***). To assess the presence of DG, we performed immunostaining of quadriceps muscles from 12-week-old M-*Dag1* KO mice, which showed the absence of DG-positive fibers (***Figure 5-figure supplement 1***). Immunoblot analysis showed that matriglycan and α-DG derived from skeletal muscle were not observed in M-*Dag1* KO mice (***Figure 5-figure supplement 1***). This is consistent with prior reports showing that only peripheral-nerve derived matriglycan of 110 kDa is observed in M-*Dag1* KO mice in the presence of EDTA, which improves the extraction of matriglycan positive α-DG and acts as a protease inhibitor by chelating calcium (***Saito et al., 2003***).

**Figure 5.**
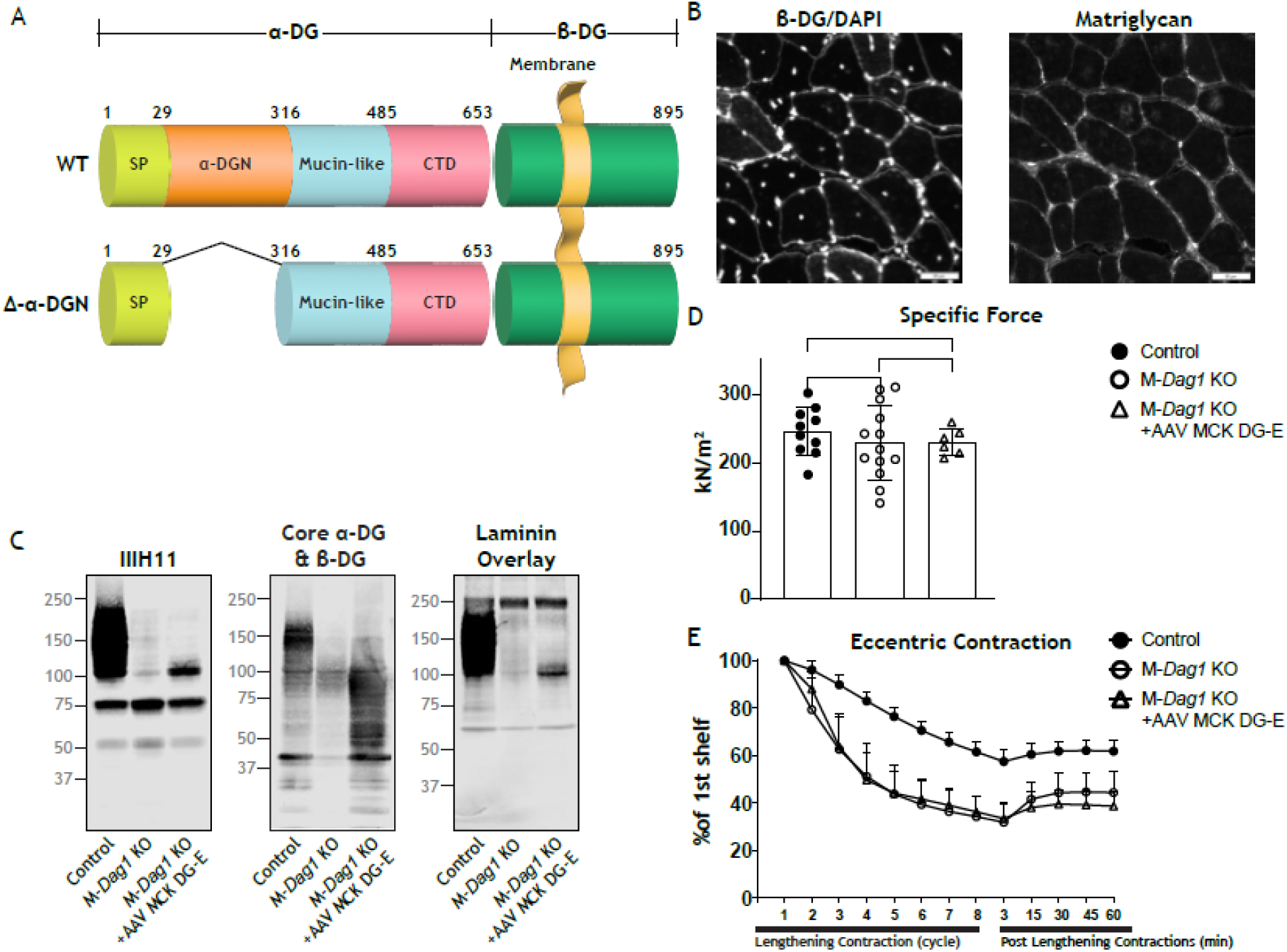
Exogenous α-DGN-deficient DG also produces short matriglycans similar to M-*Dag1* KO mice. **(A)** Schematic representation of WT DG, adeno-associated virus (AAV), and a mutant DG in which the N-terminal domain has been deleted (DG-E) adeno-associated virus. α-DG is composed of a signal peptide (SP, amino acids 1–29), an N-terminal domain (amino acids 30–316), a mucin-like domain (amino acids 317–485), and a C-terminal domain (amino acids 486–653). The green box represents β-DG. **(B)** Immunofluorescence analyses of quadriceps muscles from 12-week-old M-*Dag1* KO mice injected with AAV-MCK DG-E to detect β-DG, nuclei (DAPI) and matriglycan (IIH6). (**C**) Immunoblot analysis of skeletal muscle obtained from littermate controls (control), M-*Dag1* KO mice or M-*Dag1* KO mice injected with AAV-MCK DG-E. Glycoproteins were enriched from quadriceps skeletal muscles using WGA-agarose with 10 mM EDTA. Immunoblotting was performed to detect matriglycan (IIIH11), core α-DG and β-DG (AF6868), and laminin (overlay). **(D)** Production of specific force in EDL muscles from 12- to 17-week-old male & female M-*Dag1* KO mice (controls; closed circles, n=10); M-α-DGN KO mice (open circles, n=13); and M-*Dag1* KO+AAVMCK DG-E mice (open triangles, n=6). P-values determined by Student’s unpaired t-test; controls vs M-*Dag1* KO: p=0.4158; controls vs M-*Dag1* KO+AAVMCK DG-E: p=0.3632; M-*Dag1* KO vs M-*Dag1* KO+AAVMCK DG-E: p=0.948. **(E)** Force deficits and recovery in EDL muscles from mice in D. There is no significant difference in M-*Dag1* KO vs M-*Dag1* KO+AAVMCK DG-E as determined by Student’s unpaired t-test at any given lengthening contraction cycle or post-lengthening contraction.

We next generated an adeno-associated virus (AAV) construct of DG lacking the α-DGN (AAV-MCK DG-E; ***Figure 5A***), which we injected into M-*Dag1* KO mice through the retro-orbital sinus. A previous report found that matriglycan was not produced when a similar adenovirus construct of DG lacking the α-DGN was used to infect ES cells (***Kanagawa et al., 2004***). However, we found that matriglycan of similar size was produced in M-*Dag1* KO mice injected with AAV-MCK DG-E as in M-α-DGN KO mice (***Figure 5***). Immunofluorescence analysis of quadriceps muscle from M-*Dag1* KO mice injected with AAV-MCK DG-E showed decreased immunoreactivity to matriglycan-positive muscle fibers but restored expression of β-DG (***Figure 5B***). Immunoblot analysis of skeletal muscle from M-*Dag1* KO mice injected with AAV-MCK-DG-E showed expression of α-DG containing matriglycan around ~100-125 kDa (***Figure 5C***), which was the same size as α-DG with matriglycan in M-α-DGN KO (***Figure 2C***). The molecular weight of α-DG was decreased in muscle from these mice, similar to that observed in M-α-DGN KO mice, whereas the molecular weight of β-DG was unchanged relative to M-α-DGN KO mice (***Figure 2C***). We also observed laminin-binding at ~100-125 kDa in muscle from M-*Dag1* KO + AAV-MCK DG-E mice (***Figure 5C***). In addition, we assessed the physiologic effects of expressing DG without the N-terminal domain. We observed that the specific force was comparable in M-*Dag1* KO + AAV-MCK DG-E and M-α-DGN KO mice (***Figure 5D***) and that the two groups exhibited similar amounts of lengthening contraction-induced force decline (***Figure 5E***). Therefore, these data demonstrate that AAV-mediated delivery of exogenous DG lacking the α-DGN into an M-*Dag1* KO mouse also produces short matriglycan and exhibits the same muscle function as M-α-DGN KO mice.

Our studies show that DG lacking the α-DGN expresses a short form of matriglycan; this suggests that α-DGN is necessary for the production of full-length matriglycan. To test this hypothesis, we determined if matriglycan expression could be restored in mice lacking α-DGN. We injected M-α-DGN KO mice with an AAV expressing α-DGN (AAV-CMV α-DGN) and harvested the skeletal muscles of these mice eight to ten weeks after injection. H&E staining in M-α-DGN KO mice injected with AAV-CMV α-DGN was unchanged from M-α-DGN KO mice (***Figure 2B, 6A***). Quadriceps muscles from M-α-DGN KO mice injected with AAV-CMVα-DGN showed a reduced intensity of matriglycan relative to littermate controls (***Figure 6A***). Immunoblot analysis of these mice showed that matriglycan had a molecular weight of ~100-125 kDa and the size of α-DG was shifted down, whereas β-DG remained unchanged (***Figure 6B***). Laminin-binding was observed at ~100-125 kDa in M-α-DGN KO skeletal muscle infected with AAV-CMVα-DGN (***Figure 6B***). Collectively, this phenotype is similar to that observed in the skeletal muscles of M-α-DGN KO mice. Expressing α-DGN in M-α-DGN KO mice did not alter specific force or improve force deficits induced by lengthening contractions (***Figure 6C, 6D***). Thus, supplementing M-α-DGN KO skeletal muscle with α-DGN fails to improve matriglycan elongation.

**Figure 6.**
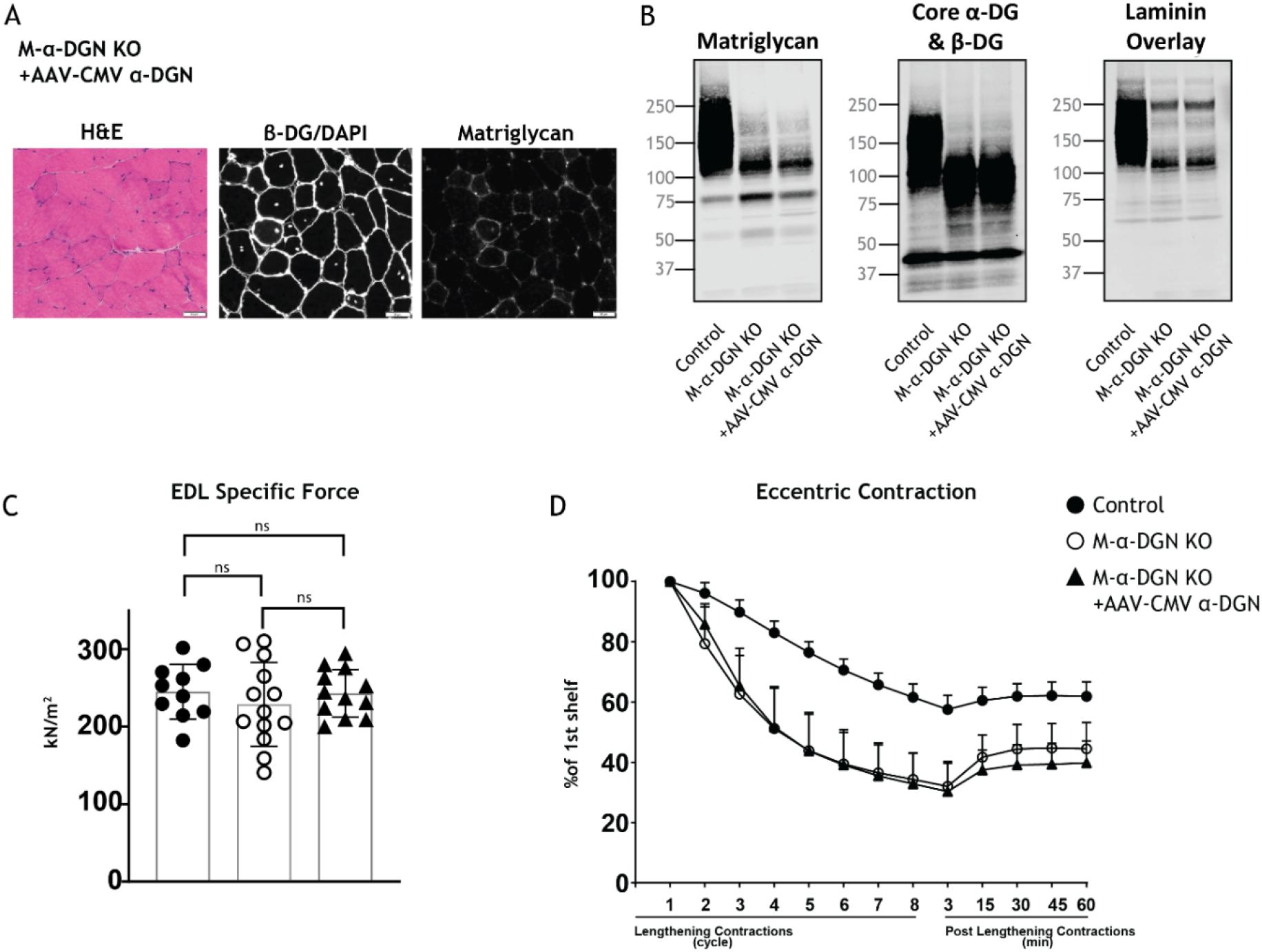
Expression of α-DGN in M-α-DGN KO mice does not rescue matriglycan elongation. **(A)** Representative sections of quadriceps muscles from 17-week-old M-α-DGN KO mice injected with AAV-CMV α-DGN. Sections were stained with H&E and immunofluorescence to detect matriglycan (IIH6) and β-DG (AP83). **(B)** Immunoblot analysis of skeletal muscle obtained from littermate controls or M-α-DGN KO mice and M-\α-DGN KO mice injected with AAV-CMV α-DGN (M-α-DGN KO+AAV-CMV α-DGN). Glycoproteins were enriched using WGA-agarose with 10 mM EDTA. Immunoblotting was performed to detect matriglycan (IIIH11), core α-DG and β-DG (AF6868), and laminin overlay. **(C)** Production of specific force in EDL muscles from 12- to 17-week-old male & female M-α-DGN WT littermates (controls; closed circles, n=10); M-α-DGN KO (open circles, n=13); and M-α-DGN KO+AAV-CMV α-DGN (closed triangles, n=12). P-values determined by Student’s unpaired t-test; controls vs M-α-DGN KO+AAV-CMV α-DGN: p=0.8759; controls vs M-α-DGN KO: p=0.4333; M-α-DGN KO vs M-α-DGN KO+AAV-CMV α-DGN: p=0.4333. **(D)** Force deficit and force recovery after lengthening contractions in EDL muscles from 12- to 17-week-old male & female M-α-DGN KO WT littermates (controls, closed circles; n=6) and M-α-DGN KO (KO, open circles; n=7) mice, and in M-α-DGN KO mice injected with AAV-CMV α-DGN (KO+AAV-CMV α-DGN, closed triangles; n=8). There is no significant difference in M-α-DGN KO vs M-α-DGN KO+AAV-CMV α-DGN as determined by Student’s unpaired t-test at any given lengthening contractions cycle or post lengthening contractions.

To determine if excess LARGE1 produces full-size matriglycan in M-α-DGN KO muscle, we evaluated immunoblot analysis of skeletal muscle from littermate controls, M-α-DGN KO, and M-α-DGN KO mice injected with AAV-MCK-Large1. M-α-DGN KO mice injected with AAV-MCK-Large1 demonstrated no change in the molecular weight of matriglycan, α-DG, and β-DG relative to M-α-DGN KO. A laminin overlay using laminin-111 also showed no change (***Figure 6-figure supplement 1***). These results indicate that even if LARGE1 is overexpressed, full-size matriglycan cannot be produced without α-DGN.

The length of matriglycan is correlated with its ability to bind ECM ligands (***Goddeeris et al., 2013***). Therefore, we hypothesized that the susceptibility to force decline by lengthening contractions would differ depending on the length of matriglycan. To test this, we performed physiological muscle tests in three different mouse models to determine the difference in susceptibility to lengthening contraction-induced reduction in force. Specifically, we used: 1) M-α-DGN KO mice, which express a short form of matriglycan, 2) *Dag1^T190M^* mice, which harbor a knock-in mutation (T190M) in *DAG1* that inhibits the DG-LARGE1 interaction and leads to incomplete post-translational modification of α-DG (***Hara et al., 2011***), and 3) C57BL/6J WT (C57) mice, which have full-length matriglycan. The percent deficit value of the 8th eccentric contraction (EC) shows the largest difference in the EC protocol; therefore, we compared these values between our three different mouse models (***Figure 7A***). M-α-DGN KO mice showed a significantly higher percent deficit (70.2% ± 5.7) compared to C57 (41.7% ± 8.0) and *Dag1^T190M^* (41.6 ± 6.7) mice, with no difference observed between the latter groups. Immunoblot analysis of laminin in skeletal muscle showed α-DG laminin-binding at ~150-250 kDa in skeletal muscle from C57 mice, ~100-150 kDa in skeletal muscle from *Dag1^T190M^* mice, and ~100-125 kDa in skeletal muscle from M-α-DGN KO mice (***Figure 7C***). Moreover, the percentage of centrally nucleated fibers differed significantly in *Dag1^T190M^* (1.73%±0.31) and M-α-DGN KO (9.28%±2.41) mice compared to C57 mice (1.22%±0.15) (***Figure 7B***). The reduction of laminin-binding activity of α-DG is thought to be the main cause of dystroglycanopathy (***Kanagawa et al., 2009; Goddeeris et al., 2013***). Indeed, we observed a reduced binding capacity (relative B_max_) for laminin-111 in solid-phase binding analyses in skeletal muscle from M-α-DGN KO and *Dag1^T190M^* mice compared to skeletal muscle from C57 mice (10.7-fold and 2.3-fold difference relative to WT, respectively) (***Figure 7D***). However, the binding capacity of skeletal muscle from *Dag1^T190M^* and M-α-DGN KO mice was higher than that of *Large^myd^* muscle (***Figure 7E***). M-α-DGN KO and *Dag1^T190M^* also displayed an increase in dissociation constant (***Figure 7E***). Collectively, these results suggest that α-DG positive matriglycan of at least ~150 kDa is sufficient to prevent force decline from lengthening contractions and significant dystrophic changes, despite a 45% reduction in laminin-binding activity.

**Figure 7.**
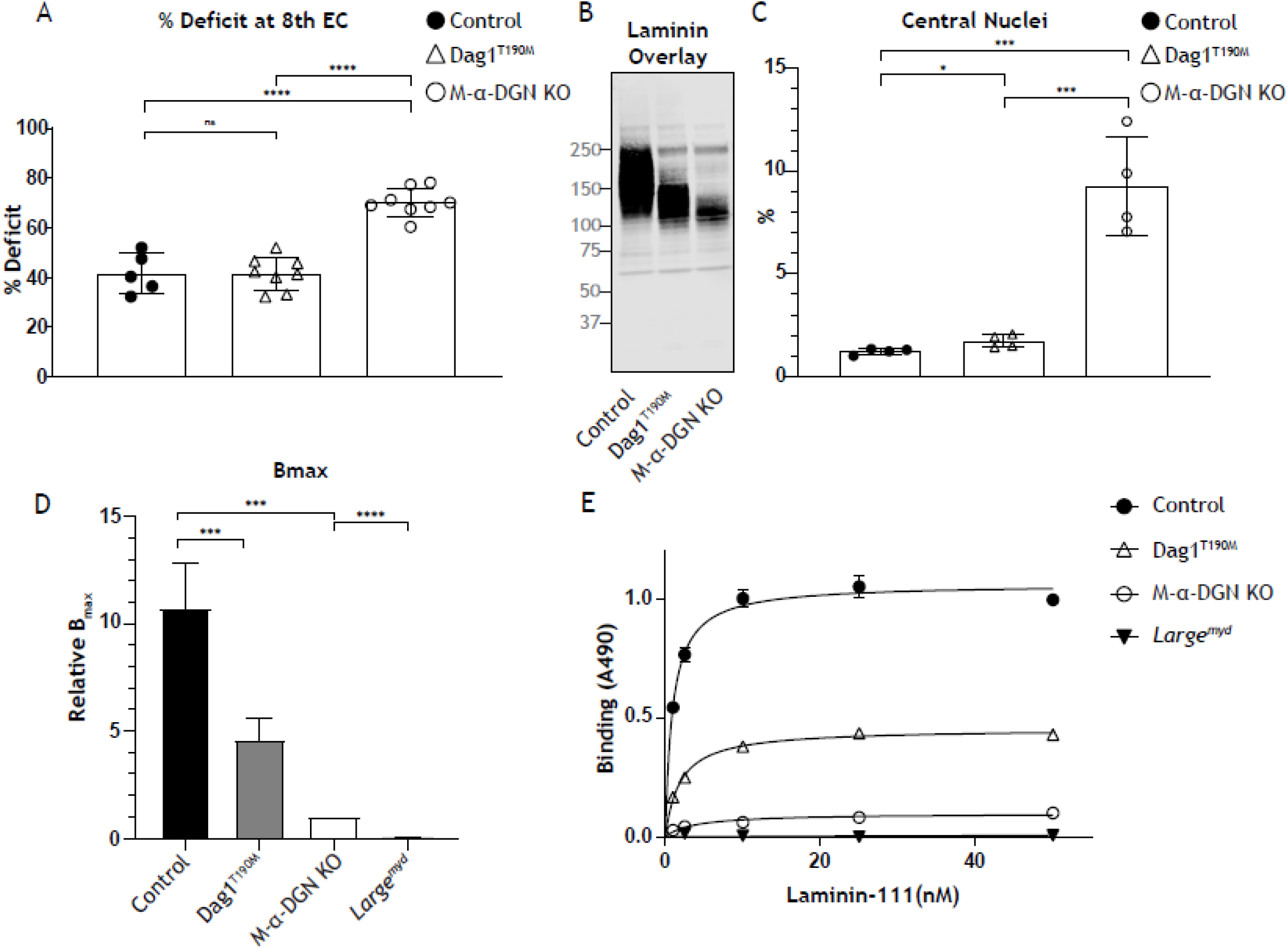
Relationship between matriglycan length and dystrophic phenotype. **(A)** % deficit of 8^th^ eccentric contraction (EC) in EDL muscles from C57BL/6J WT (control), M-α-DGN KO, and *Dag1^T190M^* mice. p-values determined by Student’s unpaired t-test; control vs *Dag1^T190M^:* p=0.0263; control and *Dag1^T190M^* vs M-α-DGN KO: p<0.001. **(B)** Percentage of muscle fibers with central nuclei in 12- to 19-week-old control, *Dag1^T190M^* and M-α-DGN KO mice; n=4 for all groups. P-values determined by Student’s unpaired t-test; control and *Dag1^T190M^* vs M-α-DGN KO: p<0.001; control vs *Dag1^T190M^:* p=0.0263. **(C)** Immunoblot analysis of quadriceps skeletal muscles from control, *Dag1^T190M^* and M-α-DGN KO mice. Glycoproteins were enriched using WGA-agarose with 10 mM EDTA. Immunoblotting was performed with laminin (laminin overlay). **(D)** Comparison of average solid-phase determined relative B_max_ values for laminin. Bmax values for M-α-DGN KO were set to 1 to allow for direct comparisons; error bars indicate s.e.m. P-values determined using Student’s unpaired t-test; control vs *Dag1^T190M^* and control vs M-α-DGN KO: p<0.01, and M-α-DGN KO vs *Large^myd^:* p<0.001. **(E)** Solid-phase analysis of laminin-binding using Laminin-111 in skeletal muscle from control, *Dag1^T190M^*, M-α-DGN KO, and *Large^myd^* KO mice (three replicates for each group). Control K_d_: 0.9664 ± 0.06897 nM; *Dag1^T190M^* K_d_: 1.902 ± 0.1994 nM; and M-α-DGN KO K_d_: 2.322 ± 0.6114 nM.

## Discussion

Functional glycosylation of α-DG requires α-DGN (***Kanagawa et al., 2004; Hara, Kanagawa, et al., 2011***). However, it remains unclear how the loss of α-DGN affects matriglycan synthesis. Here, we show that the lack of α-DGN does not preclude matriglycan synthesis entirely. Instead, in the absence of α-DGN, LARGE1 synthesizes a short, non-elongated form of matriglycan on α-DG (~ 100-125 kDa), which demonstrates that the N-terminal domain is required for matriglycan elongation. Thus, LARGE1-αDGN holds the enzyme-substrate complex together over multiple cycles of sugar addition. These findings build on our previous study demonstrating that phosphorylation of the core M3 trisaccharide by POMK is also necessary for matriglycan elongation (***Walimbe et al., 2020***). Thus, the generation of full-length mature matriglycan on α-DG (~150-250 kDa) by *LARGE1* requires both POMK and α-DGN to be bound to DG; in the absence of either, a shorter form is generated.

In our study, muscle-specific deletion of α-DGN resulted in the production of short forms of matriglycan on α-DG (~100-125 kDa). Mice lacking α-DGN exhibited low bodyweight and grip strength, and histological characterization of quadriceps muscles revealed mild muscular dystrophy and a lack of homogeneous matriglycan expression. Physiological examination revealed that M-α-DGN KO muscle was susceptible to lengthening contraction-induced force decline, although specific force was maintained. These results are consistent with those obtained when α-DGN-deleted DG was administered to muscle-specific DG KO mice and indicates that DG lacking α-DGN produces short forms of matriglycan, which does not prevent dystrophic muscle changes in this mouse model. Furthermore, DG in the postsynaptic membrane is known to play a key role in synaptic maturation (***Nishimune et al., 2008***). However, the NMJs in M-α-DGN KO mice in our study were abnormal and irregularly shaped. This indicates both that DG and matriglycan are required for synaptic maturation.

If LARGE1 binding to α-DGN enables its ability to elongate matriglycan, then we would expect that rescuing M-α-DGN KO skeletal muscle with α-DGN would restore the expression of full-length matriglycan. However, this failed to occur and indicates that solely restoring α-DGN expression is not sufficient for LARGE1 to elongate matriglycan. These results indicate that the ability for LARGE1 to elongate matriglycan requires α-DGN to be attached to DG. This finding is consistent with data showing that matriglycan is not elongated when α-DGN is deleted, even when LARGE1 is overexpressed. Therefore, α-DGN acts as a recognition site for the glycosyltransferase *LARGE1* and establishes a model where α-DGN, together with phosphorylated core M3, anchors *LARGE1* to the matriglycan production site to enable its synthesis and elongation. Notably, although the molecular recognition of α-DGN by *LARGE1* is considered essential for the expression of functional DG (***Kanagawa et al., 2004***) our results show that *LARGE1* can synthesize a short non-elongated form of matriglycan in the absence of α-DGN, indicating that LARGE1 is capable of adding matriglycan to α-DG independent of its interaction with α-DGN.

To determine how much matriglycan is needed to prevent lengthening contraction-induced reduction in force, we used mice that express different sizes of matriglycan. Muscle from M-α-DGN KO mice showed an increased force deficit and a 7.6-fold increase in centrally nucleated fibers compared to muscle from C57 mice indicating that short forms of matriglycan do not prevent dystrophic changes. However, despite the lower amount of matriglycan in muscle from *Dag1^T190M^* mice compared to that from C57 mice, the force deficit was not different between the two groups, and centrally nucleated fibers were increased in α-DGN mutant (T190M) mice by only 1.4-fold. This indicates that short matriglycan, if over 150kDa, can prevent muscular dystrophy.

Muscular dystrophy is not observed in a mouse model of Fukuyama congenital muscular dystrophy, which occurs due to a retrotransposition insertion in the mouse *fukutin* ortholog and causes laminin-binding at 50% of normal levels (***Kanagawa et al., 2009***). In *Dag1^T190M^* mice, the laminin-binding level is about 45% of normal, which likely explains a mild increase in centrally nucleated fibers compared to C57 muscle. However, in muscle from M-α-DGN KO mice, the laminin-binding level is only 9% relative to that of C57 mice and leads to a marked increase in centrally nucleated fibers and force deficit induced by lengthening contractions. This indicates that matriglycan length is critical for regulating damage induced by lengthening contractions and that the production of ~120-150 kDa α-DG significantly prevents dystrophic change, suggesting that this pathologic effect can be prevented without the expression of full-length matriglycan. Thus, our results describe a relationship between matriglycan size, damage induced by lengthening contractions, and the degree of dystrophic change. However, the difference between the abundance of central nuclei and the results of damage due to lengthening contractions in C57 and *Dag1* mice indicate that other factors likely contribute to normal physiologic function in muscle.

Collectively our study demonstrates that α-DG with α-DGN is required for the synthesis of full-length matriglycan on α-DG (~150-250 kDa). In the absence of α-DGN, LARGE1 can synthesize a short non-elongated form of matriglycan on α-DG (~100-125kDa) in skeletal muscle in a process that is independent of its interaction with α-DGN. These findings are essential for a complete understanding of the mechanisms underlying matriglycan synthesis and show that matriglycan length regulates the severity of muscular dystrophy and may serve as a therapeutic target for the treatment of α-dystroglycanopathy.

## Materials and methods

### Animals

All mice were maintained in a barrier-free, specific pathogen-free grade facility and had access to normal chow and water *ad libitum*. All animals were manipulated in biosafety cabinets and change stations using aseptic procedures. The mice were maintained in a climate-controlled environment at 25°C on a 12/12 hour light/dark cycle. Animal care, ethical usage, and procedures were approved and performed in accordance with the standards set forth by the National Institutes of Health and the University of Iowa Animal Care and Use Committee (IACUC). Mouse lines used in the study that have been previously described are: *Dag1^-/-^* (JAX# 006836; ***Williamson et al., 1997***), *Dag^flox^* (JAX# 009652; ***Cohn et al., 2002***), *Dag1^Δα-DGN^* (***de Greef et al., 2019***), *Dag1^T190M^* (***Hara et al., 2011***), *Large1^myd^* (JAX# 000300) (***Lane et al., 1976***), *Mck^cre^* (JAX# 006475) (***Brüning et al., 1998***), Pax7^cre^ (JAX# 010530) (***Keller et al., 2004***), and *Mck*^cre^ *Pax7^cre^ POMK^flox^* (***Walimbe et al., 2020***).

### Muscle-specific DG knockout mice (*Pax7^cre^ Dag1^flox/flox^*)

Male mice expressing the *Pax7-Cre* transgene were bred to female mice that were homozygous for the floxed *Dag1* allele (*Dag1^flox/flox^*). Male F1 progeny with the genotype *Pax7^Cre^; Dag1^flox/+^* were bred to female *Dag1^flox/flox^* mice. A *Cre* PCR genotyping protocol was used to genotype the *Cre* allele using standard *Cre* primers. The primers used were Sense: TGATGAGGTTCGCAAGAACC and Antisense: CCATGAGTGAACGAACCTGG. Genotyping of *Pax7_Cre_; Dag1^flox/flox^* mice was performed by Transnetyx using real-time PCR.

### Muscle-specific α-DGN knockout mice (M-α-DGN KO)

Male mice expressing the *Pax7-Cre* transgene were bred to female mice that were heterozygous for the *Dag1^Δ α-DGN^* allele (*Dag1^wt/Δα-DGN^*). Male F1 progeny with the genotype *Pax7^Cre^; Dag1^wt/Δα-DGN^* were bred to female mice homozygous for the floxed *Dag1* allele (*Dag1^flox/flox^*). Genotyping of *Pax7^Cre^Dag1^flox/Δα-DGN^* mice was performed by Transnetyx using real-time PCR. For studies with M-α-DGN KO mice, three mice of each genotype (control and *Pax7^Cre^Dag1^flox/Δα-DGN^*) were used.

Littermate controls were employed whenever possible. The number of animals required was based on previous studies (***de Greef et al., 2016; Goddeeris et al., 2013, Walimbe et al., 2020***) and experience with standard deviations of the given techniques.

### Forelimb grip strength test

Forelimb grip strength was measured at three months using previously published methods (***de Greef et al., 2016, Walimbe, et al., 2020***). A mouse grip strength meter (Columbus Instruments, Columbus, OH) was mounted horizontally, with a non-flexible grid connected to the force transducer. The mouse was allowed to grasp the grid with its two front paws and then pulled away from the grid by its tail until the grip was broken. This was done three times over five trials, with a one-minute break between each trial. The gram force was recorded per pull, and any pull where only one front limb or any hind limbs were used was discarded. If the mouse turned, the pull was also discarded. After 15 pulls (five sets of three pulls), the mean of the three highest pulls of the 15 was calculated and reported. Statistics were calculated using GraphPad Prism 8 software. Student’s t-test was used (two-sided). Differences were considered significant at a p-value less than 0.05. Graph images were also created using GraphPad Prism and the data in the present study are shown as the means + / - SD unless otherwise indicated.

### Body weight measurements

Mice were weighed as previously described (***de Greef et al., 2016, Walimbe et al., 2020***). Weights were measured after testing grip strength using a Scout SPX222 scale (OHAUS Corporation, Parsippany, NJ), and the tester was blinded to genotype. Statistics were calculated using GraphPad Prism 8 software and Student’s t-test was used (two-sided). Differences were considered significant at a p-value less than 0.05. Graph images were also created using GraphPad Prism and the data in the present study are shown as the means + / - SD unless otherwise indicated.

### Measurement of *in vitro* muscle function

To compare the contractile properties of muscles, EDL muscles were surgically removed as described previously (***Rader et al., 2016; de Greef et al., 2016, Walimbe et al., 2020***). The muscle was immediately placed in a bath containing a buffered physiological salt solution (composition in mM: NaCl, 137; KCl, 5; CaCl_2_, 2; MgSO_4_, 1; NaH_2_PO_4_, 1; NaHCO_3_, 24; glucose, 11). The bath was maintained at 25°C, and the solution was bubbled with 95% O_2_ and 5% CO_2_ to stabilize pH at 7.4. The proximal tendon was clamped to a post and the distal tendon was tied to a dual mode servomotor (Model 305C; Aurora Scientific, Aurora, ON, Canada). Optimal current and whole muscle length (L_0_) were determined by monitoring isometric twitch force. Optimal frequency and maximal isometric tetanic force (F_0_) were also determined. The muscle was then subjected to an EC protocol consisting of eight ECs at three-minute intervals. A fiber length L_f_-to-L_0_ ratio of 0.45 was used to calculate L_f_. Each EC consisted of an initial 100 millisecond isometric contraction at optimal frequency immediately followed by a stretch of L_o_ to 30% of L_f_ beyond L_o_ at a velocity of 1 L_f_/s at optimal frequency. The muscle was then passively returned to L_o_ at the same velocity. At 3, 15, 30, 45, and 60 minutes after the EC protocol, isometric tetanic force was measured. After the analysis of the contractile properties, the muscle was weighed. The CSA of muscle was determined by dividing the muscle mass by the product of L_f_ and the density of mammalian skeletal muscle (1.06 g/cm3). The specific force was determined by dividing F_o_ by the CSA (kN/mm2). 18–20-week-old male mice were used, and right and left EDL muscles from each mouse were employed whenever possible, with five to eight muscles used for each analysis. Each data point represents an individual EDL. Statistics were calculated using GraphPad Prism 8 software and Student’s unpaired t-test was used (two-sided). Differences were considered significant at a p-value less than 0.05.

### H&E and immunofluorescence analysis of skeletal muscle

Histology and immunofluorescence of mouse skeletal muscle were performed as described previously (***Goddeeris et al., 2013***). Mice were euthanized by cervical dislocation and directly after sacrifice, quadriceps muscles were isolated, embedded in OCT compound and then snap frozen in liquid nitrogen-cooled 2-methylbutane. 10 μM sections were cut with a cryostat (Leica CM3050S Research Cryostat; Amsterdam, the Netherlands) and H&E stained using conventional methods. Whole digital images of H&E-stained sections were taken by a VS120-S5-FL Olympus slide scanner microscope (Olympus Corporation, Tokyo, Japan). For immunofluorescence analyses, a mouse monoclonal antibody to matriglycan on α-DG (IIH6, 1:100 dilution, Developmental Studies Hybridoma Bank, University of Iowa; RRID:AB_2617216) was added to sections overnight at 4 °C followed by Alexa Fluor-conjugated goat IgG against mouse IgM (Invitrogen, Carlsbad, CA, 1:500 dilution) for 40 minutes. The sections were also stained with rabbit polyclonal antibody to β-DG (AP83; 1:50 dilution) followed by Alexa Fluor-conjugated 488 Goat anti-rabbit IgG (1:500). Whole sections were imaged with a VS120-S5-FL Olympus slide scanner microscope. Antibody IIH6 is a mouse monoclonal to matriglycan on α-DG (***Ervasti and Campbell, 1991***), and AP83 is a rabbit polyclonal antibody to the C-terminus of β-DG (***Ervasti and Campbell, 1991***), both of which have been described previously.

For histologic analysis of skeletal muscle, H&E staining on 10 μM frozen section was performed using the Leica ST5020 Multistainer workstation (Leica Biosystems, Buffalo Grove, IL) according to the manufacturer’s instructions. For immunofluorescence analysis, unfixed frozen serial sections (7 μM) were incubated with primary antibodies for one hour, and then with the appropriate biotinylated secondary antibodies for 30 minutes followed by streptavidin conjugated to Alexa Fluor 594 (ThermoFisher Scientific, UK) for 15 minutes. Primary antibodies used were mouse monoclonal: α-DG IIH6 (clone IIH6C4) (***Ervasti and Campbell, 1991***), β-DG (Leica, Milton Keynes, UK; clone 43DAG1/8D5). All washes were made in PBS and incubations were performed at room temperature. Sections were evaluated with a Leica DMR microscope interfaced to MetaMorph (Molecular Devices, Sunnyvale, CA).

### Neuromuscular Junction (NMJ) Morphology

Immediately upon harvest, EDL muscles were washed in PBS three times for five minutes each. EDL muscles were fixed in 4% paraformaldehyde for 20 minutes followed by three washes in PBS. Fixed muscle samples were split into three to four fiber bundles before incubating in 3% Triton-X 100/PBS for three hours at 4 °C. Muscles were subsequently washed in PBS followed by blocking at 4 °C for four hours in Background Buster (Innovex; NB306). Samples incubated with primary antibodies against neurofilament H (NF-H; EnCor; CPCA-NF-H) at 1:1,000 and synaptophysin (Thermo Fisher Scientific; MA5-14532) at 1:100 diluted in 5% Background Buster/1% Triton-X 100/PBS at 4 °C overnight. The muscles were then washed with PBS and incubated with fluorescently conjugated secondary antibodies and Alexa Fluor 488-conjugated α-bungarotoxin (Invitrogen; B13422) diluted in 5% Background Buster/PBS for two hours. Images were acquired using an Olympus FLUOVIEW FV3000 confocal laser scanning microscope. Complete *enface* NMJs were identified and acquired with Z-stacks using 60x and 100x objectives. Maximum intensity Z-stacks were reconstructed with the FV31S (Olympus) software and deconvoluted with cellSens Dimension (Olympus). Blinded observers analyzed α-BTX-488-labeled AChR cluster formations to determine irregularities, fragmentation, synaptic size, and dispersion. Irregularities included AChR plaques, AChR perforated plaques, ring-shaped or c-shaped clusters, and extensive fragmentation. Fragmentation was determined by the number of identifiable individual AChR clusters within the footprint of the synapse. FUJI ImageJ software was used for semi-automatic analysis of AChR clusters. Synaptic size refers to the total perimeter or footprint of the postsynapse. AChR cluster dispersion was determined by the (total stained area/total area) *100.

### Tissue biochemical analysis

Mouse skeletal muscle was minced into small pieces and homogenized with polytron (Kinematica, PT10-35) three times for 10 seconds at power 4 to 5 in 15 ml of TBS (150 mM NaCl) with 1% TX100 and 10 mM EDTA, and protease inhibitors (per 10 mL buffer: 67 mL each of 0.2 M phenylmethylsulfonylfluoride (PMSF), 0.1 M benzamidine, and 5 μL of each of leupeptin (Sigma/Millipore) 5 mg/mL, pepstatin A (Millipore) 1 mg/mL in methanol, and aprotinin (Sigma-Aldrich) 5 mg/mL. The samples were incubated in a cold room 1 hr. with rotation. The samples were centrifuged in a Beckman Coulter Avanti J-E centrifuge for 30 minutes at 20,000xg, 4 °C. The supernatant was combined with WGA slurry at 600 μL per gram of starting muscle and rotated at 4C over night.

The WGA beads were washed using 10X volume of WGA beads/wash 3X for three minutes at 1000 x g with 0.1%Tx/TBS, plus protease inhibitors. After the final wash, the WGA beads (Vector Laboratories, AL-1023) were eluted with Laemmli Sample Buffer (LSB) at 600 μL per gram of starting material at 99 °C for 10 minutes. The final concentration was 1.11 mg skm/μL beads and LSB. Samples were loaded (beads and LSB) in a 3-15% gradient gel. The proteins were transferred to PVDF-FL membranes (Millipore) as previously published (***Michele et al., 2002; Goddeeris et al., 2013***). EDTA (10 mM) was used in the homogenization to more efficiently extract α-DG containing matriglycan in the muscle homogenates. (***Figure 7-figure supplement 1 WT and POMK***), while EDTA had no effect on *Large^myd^* α-DG (matriglycan-negative) extraction (***Figure 7-figure supplement 1 Large^myd^***).

### Immunoblotting and ligand overlay

The mouse monoclonal antibody against matriglycan on α-DG (IIH6, Developmental Studies Hybridoma Bank, University of Iowa; RRID:AB_2617216) was characterized previously and used at 1:100 (***Ervasti and Campbell, 1991***). The polyclonal antibody, AF6868 (R&D Systems, Minneapolis, MN; RRID:AB_10891298), was used at a concentration of 1:100 for immunoblotting the core α-DG and β-DG proteins, and the secondary was a donkey anti-sheep (LI-COR Bioscience, Lincoln, NE) used at 1:10,000 concentration. The mouse xxxx antibody against matriglycan on α-DG (III HII) was previously used (***Groh et al., 2009***). Blots were developed with infrared (IR) dye-conjugated secondary antibodies (***Walimbe et al., 2020)*** and scanned using the Odyssey infrared imaging system (LI-COR Bioscience). Blot images were captured using the included Odyssey image-analysis software.

Laminin overlay assays were performed as previously described (***Michele et al., 2002; Goddeeris et al., 2013***). Immobilon-FL membranes were blocked in laminin-binding buffer (LBB: 10 mM triethanolamine, 140 mM NaCl, 1 mM MgCl_2_, 1 mM CaCl_2_, pH 7.6) containing 5% milk followed by incubation with mouse Engelbreth-Holm-Swarm (EHS) laminin (ThermoFisher, 23017015) overnight at a concentration of 7.5 nM at 4 °C in LBB containing 3% bovine serum albumin (BSA) and 2 mM CaCl_2_. Membranes were washed and incubated with anti-laminin antibody (L9393; Sigma-Aldrich 1:1000 dilution) followed by IRDye 800 CW dye-conjugated donkey anti-rabbit IgG (LI-COR, 926–32213) at 1:10,000.

### Digestion of α-DG with exoglycosidases

β-glucuronidase from *Thermotoga maritima* and α-xylosidase from *Sulfolobus solfataricus* were cloned into pET-28a (+) vector between NheI/XhoI sites in frame with the N-terminal 6xHis tag. The plasmids (20ng each) were chemically transformed into 50μl BL21DE3 One shot competent cells. One colony each was picked and inoculated in 20ml LB (with kanamycin 50 μg/ml) overnight at 37 °C. The next day, 10 ml of the overnight culture was inoculated into 1-liter LB (with kanamycin 50μg/ml). After reaching 0.6 OD at 600 nm the cultures were induced with 1 mM IPTG and incubated at 16 °C overnight. The next day the cells were centrifuged at 5000g, for 10 minutes at 4 °C. Cell pellets were stored at −80 °C until ready for purification.

The pellets were dissolved in 20 ml homogenization buffer (50 mM Tris-Cl, 150 mM NaCl, 1% TX-100, and all protease inhibitors) per liter culture. The cells were stored again overnight in 50 ml falcon tubes at −80 °C for ice crystal formation. Cells were thawed the next day for purification. Nuclease (Pierce) was added at 1.25kU and cells were sonicated at power level four-five for four times with 10s intervals in between at 4 °C. Cells were then centrifuged at 15000g for 20 minutes at 4 °C. The supernatant was heat fractionated at 75 °C for 10 minutes after which it was centrifuged at 15000g for 30 minutes at 4 °C. Meanwhile, a TALON superflow metal affinity column was prepared by packing 3 ml of resin and equilibrating with wash buffer 1 (50 mM Tris-Cl, 100 mM NaCl, 0.1% TX-100, all PIs). All further purification steps were performed at 4 °C. The extract was applied to the column three times, such that each time, the extract was incubated with the column for 15-30 minutes on gentle rocking platform. All flowthrough was saved. The column was washed three times with wash buffer 1. All washes were saved. The column was next washed with high salt wash buffer (50 mM Tris-Cl, 500 mM NaCl, 0.1% TX-100, all PIs) to remove nonspecific interactions and the high salt wash was saved. Proteins were then eluted with elution buffer (50 mM Tris-Cl, 100 mM NaCl, 0.1% TX-100 and 300 mM Imidazole) in five fractions of three milliliters each. The relevant fractions (elute 1 and 2) were pooled together, and buffer exchanged with 1XPBS pH 7.4 with 30 kDa concentrators (Amicon). 100 μl was loaded on SDS PAGE from all fractions and washes to visualize with Coomassie.

WGA enriched glycoproteins (elutes) were buffer exchanged with Sodium acetate buffer pH 5.5 using 30 kDa concentrators and heated for five minutes in the presence of 10 mM β-mercaptoethanol at 99 °C. All protease inhibitors were added after the mixture cooled down. 50 μl of each enzyme was added per 500 μl of WGA-enriched and buffer-exchanged glycoproteins. The initial time point was aliquoted as T_o_ and the rest was incubated at 75 °C with 600 rpm shaking for 16 hours.

### AAV vector production and AAV injection

The sequence encoding mouse *like-acetylglucosaminyltransferase-1* (*Large1*) was synthesized (Genscript, Piscataway, NJ) and cloned into the AAV backbone under the transcriptional control of the ubiquitous CMV promoter. The AAV2/9 vector contains the genome of serotype 2 packaged in the capsid from serotype 9 and was selected due to its ability to improve muscle transduction efficiency as well as alter tropism. The vector AAV2/9-CMV-*Large1* was generated by the University of Iowa Viral Vector Core Facility. For adult mice, 100 μL (4.35 x 10^12^ vg) of the vector solution was administered once intraperitoneally or intravenously via the retro-orbital (RO) sinus. The sequence encoding mouse *like-acetylglucosaminyltransferase-1* (*Large1*) was synthesized (Genscript, Piscataway, NJ) and cloned into the AAV backbone under the transcriptional control of the muscle-specific MCK promoter (gift from Jeff Chamberlain). The vector AAV2/9-MCK-*Large1* was generated by the University of Iowa Viral Vector Core Facility. For adult mice, 100 μL (2.55 x 10^12^ vg) of the vector solution were administered once intraperitoneally or intravenously via the retro-orbital (RO) sinus. The sequence encoding mouse *α-DG lacking the N-terminal domain* (*H30 – A316*) was synthesized (Genscript) and cloned into the AAV backbone under the transcriptional control of the muscle-specific MCK promoter. The vector AAV2/9-MCK-*DG-E* was generated by the University of Iowa Viral Vector Core Facility. For adult mice, 100 μL (6.17 x 10^11^ vg) of the vector solution was administered once intraperitoneally or intravenously via the retro-orbital (RO) sinus. The sequence encoding mouse *alpha-DG N terminal domain(α-DGN*) was synthesized (Genscript) and cloned into the AAV backbone under the transcriptional control of the ubiquitous CMV promoter. The AAV2/9 vector contains the genome of serotype 2 packaged in the capsid from serotype 9 and was selected due to its ability to improve muscle transduction efficiency as well as alter tropism. The vector AAV2/9CMV*α-DGN* was generated by the University of Iowa Viral Vector Core Facility. For adult mice, 100 μL (1.7 x 10^12^ vg) of the vector solution was administered once intraperitoneally or intravenously via the retro-orbital (RO) sinus.

### Solid-phase assay

Solid-phase assays were performed as described previously (***Michele et al., 2002; Goddeeris et al., 2013***). Briefly, WGA N-acetyl-glycosamine buffer eluates were diluted 1:50 in TBS and coated on polystyrene ELISA microplates (Costar 3590) overnight at 4 °C. Plates were washed in LBB and blocked for two hours in 3% BSA/LBB at room temperature. The wells were washed with 1% BSA/LBB and incubated for one hour with L9393 (1:5000 dilution) in 3% BSA/LBB followed by incubation with Horseradish Peroxidase (HRP)-conjugated anti-rabbit IgG (Invitrogen, 1:5000 dilution) in 3% BSA/LBB for 30 minutes. Plates were developed with o-phenylenediamine dihydrochloride and H_2_O_2_, and reactions were stopped with 2 N H_2_SO_4_. Absorbance per well was read at 490 nm by a microplate reader.

### Statistics

The included Shimadzu post-run software was used to analyze LARGE1 activity in mouse skeletal muscle, and the percent conversion to the product was recorded. The means of three experimental replicates (biological replicates, where each replicate represents a different pair of tissue culture plates or animals, i.e. control and knockout) were calculated using Microsoft Excel, and the mean percent conversion to product for the WT or control sample (control mouse skeletal muscle or *M-α-DGN KO* mouse skeletal muscle and *Large^myd^* mouse skeletal muscle, respectively) reaction was set to one. The percent conversion of each experimental reaction was subsequently normalized to that of the control, and statistics on normalized values were performed using GraphPad Prism 8. For analysis of LARGE1 activity in mouse skeletal muscle, Student’s t-test was used (two-sided). Differences were considered significant at a p-value less than 0.05. Graph images were also created using GraphPad Prism and the data in the present study are shown as the means + / - SD unless otherwise indicated. The number of sampled units, n, upon which we report statistics for *in vivo* data, is the single mouse (one mouse is n = 1).

### Data Availability

All data generated or analyzed during this study are included in this published article.

## Acknowledgements

We thank Keith Garringer for technical assistance and the University of Iowa Viral Vector Core for generating the adeno-associated viral vector (http://www.medicine.uiowa.edu/vectorcore). The MCK promoter was a gift from Jeff Chamberlain (University of Washington, Seattle, WA). We are grateful to Dr. Jennifer Barr of the Scientific Editing and Research Communication Core at the University of Iowa Carver College of Medicine for her critical reading of the manuscript. We are also grateful to Amber Mower for her assistance with administrative support and Rachel Poe for her support in figure design.

## Ethics

Animal experimentation: This study was performed in strict accordance with the recommendations in the Guide for the Care and Use of Laboratory Animals of the National Institutes of Health. All animal experiments were approved by the Institutional Animal Care and Use Committee (IACUC) protocols of the University of Iowa (#0081122).

## Author contributions

Conceptualization: H.O., K.P.C.

Methodology: H.O., J.M.H., I.C., M.E.A., D.V., A.S.W., S.J., Y.H., F.S., K.M., K.P.C.

Formal analysis: H.O., J.M.H., I.C., M.E.A., D.V., A.S.W., S.J., K.P.C.

Investigation: H.O., J.M.H., I.C., M.E.A., D.V., A.S.W., S.J., Y.H., F.S., K.M., K.P.C.

Writing – original draft preparation: H.O., K.P.C.

Writing – review & editing: H.O., J.M.H., I.C., M.E.A., D.V., A.S.W., S.J., K.P.C.

Supervision: K.P.C.

Project administration: K.P.C.

Funding acquisition: K.P.C.

---

**Figure 2-figure supplement 1.**
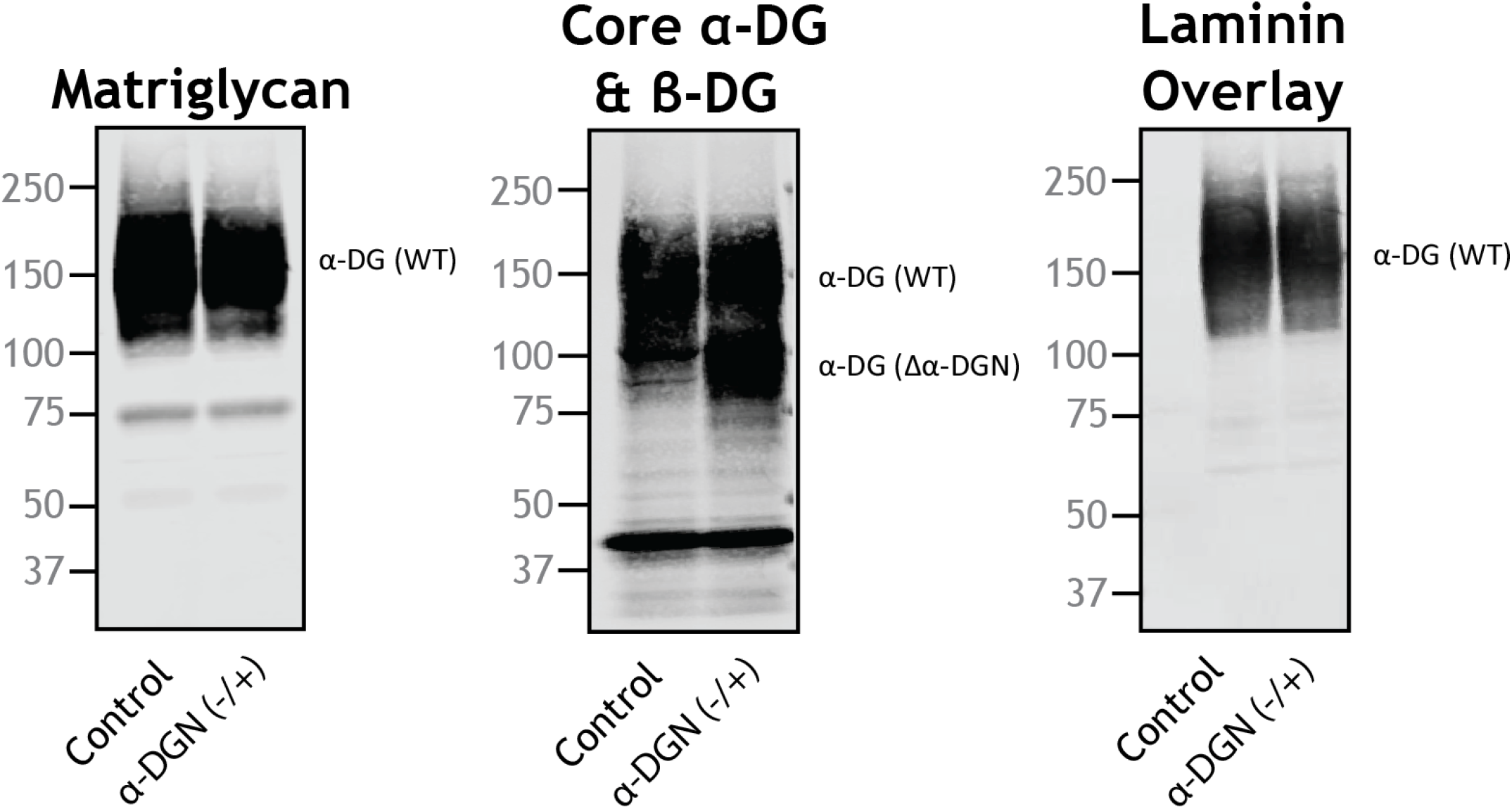
Heterozygous mice (+/-) for constitutive deletion of α-DGN have two different sizes of α-DG. Immunoblot analysis of skeletal muscle from littermate controls or mice that were heterozygous for the α-DGN KO allele (α-DGN (-/+)). Glycoproteins were enriched from quadriceps skeletal muscles of mice using WGA-agarose with 10 mM EDTA. Immunoblotting was performed to detect matriglycan (IIIH11), core α-DG and β-DG (AF6868), and laminin overlay. α-DG in WT control muscle (α-DG(WT)) and α-DG in α-DGN-deficient muscle (α-DG(Δα-DGN)) are indicated on the right. Molecular weight standards in kilodaltons (kDa) are shown on the left.

**Figure 2-figure supplement 2.**
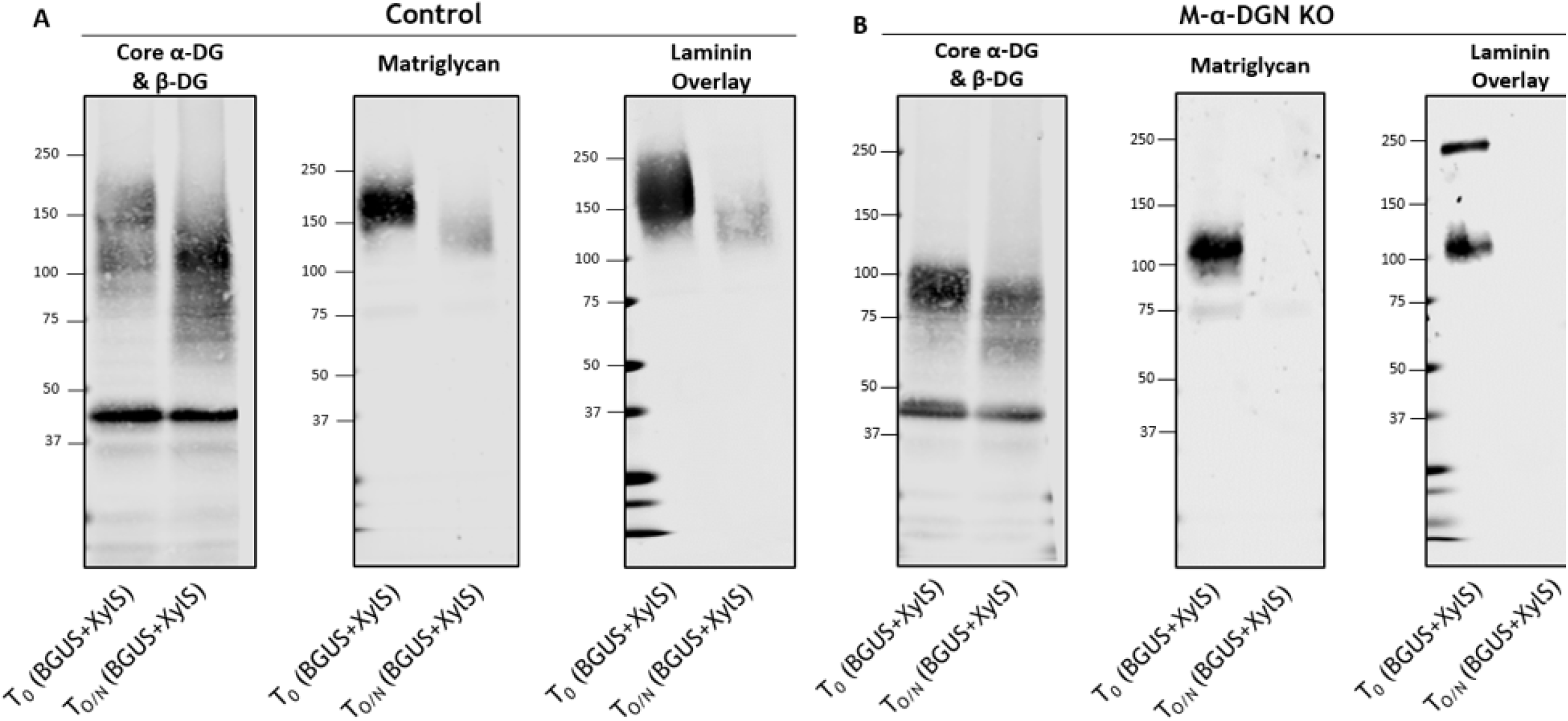
The short 100-120kDa band in M-α-DGN KO is matriglycan. **(A)** Immunoblot analysis of total skeletal muscle from control mice after digestion with enzymes β-glucuronidase and α-xylosidase. Glycoproteins were enriched using wheat-germ agglutinin (WGA)-agarose with 10 mM EDTA and incubated overnight with β-glucuronidase (BGUS) and α-xylosidase (XyIS). Immunoblotting was performed to detect matriglycan (IIH6), core α-DG and β-DG (AF6868), and laminin overlay before (T_o_) and after overnight digestion (T_O/N_). **(B)** Immunoblot analysis of M-α-DGN KO total skeletal muscle after digestion with enzymes β-glucuronidase and α-xylosidase. Glycoproteins were enriched using wheat-germ agglutinin (WGA)-agarose with 10 mM EDTA and incubated overnight with β-glucuronidase and α-xylosidase. Immunoblotting was performed to detect matriglycan (IIH6), core α-DG and β-DG (AF6868), and laminin overlay before (T_o_) and after digestion (T_O/N_). Molecular weight standards in kilodaltons (kDa) are shown on the left.

**Figure 4-figure supplement 1.**
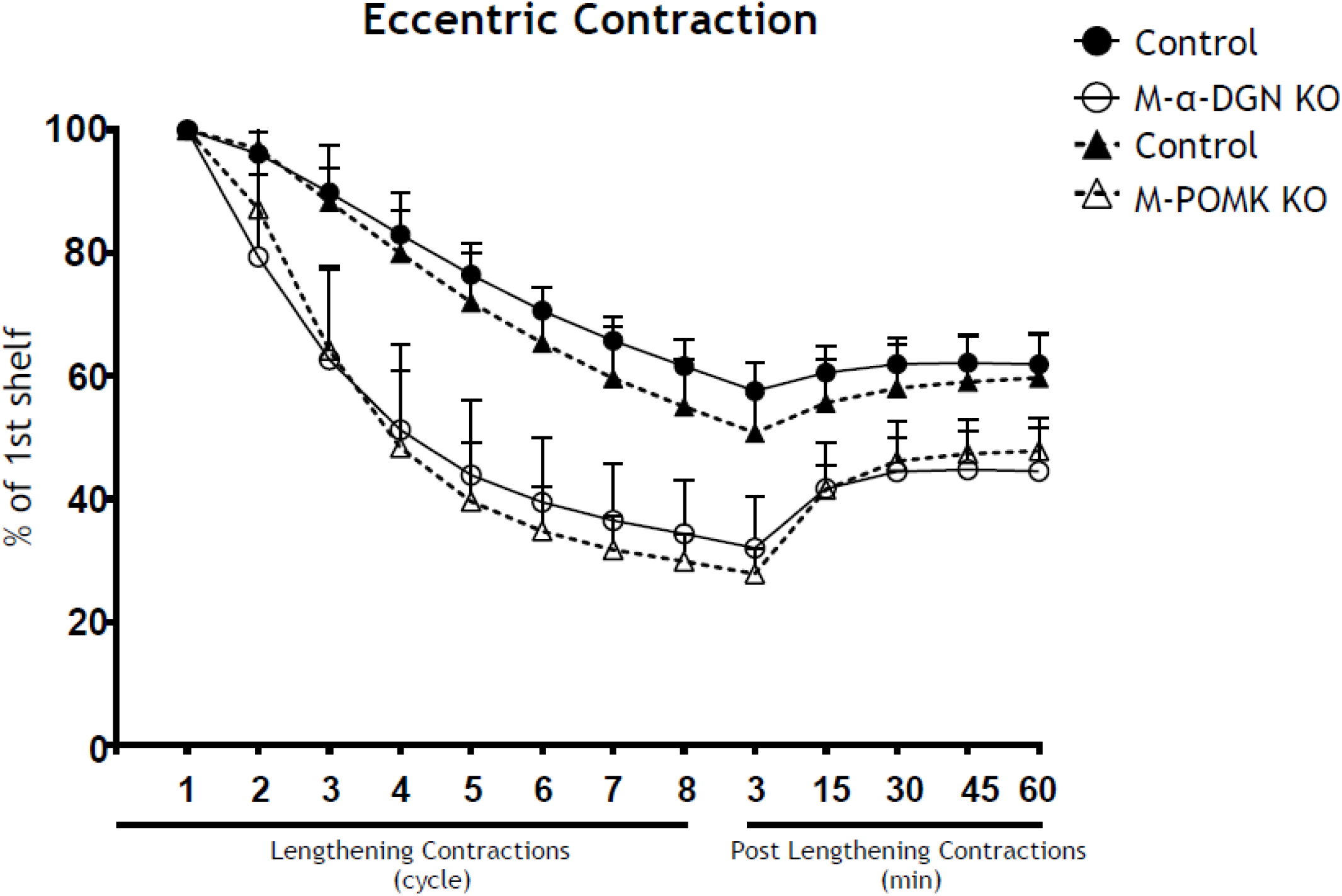
α-DGN-deficient muscle and POMK-deficient muscle with similar short forms of matriglycan exhibit similar lengthening contraction-induced force decline. Force deficit and force recovery after eccentric contractions in EDL muscles from 12- to 17-week-old male & female controls (closed circles; n=7), M-α-DGN KO (open circles; n=7), M-POMK littermate controls (closed triangles; n=3), and M-POMK KO (open triangles; n=4) mice. There is no significant difference in M-α-DGN KO vs M-POMK KO as determined by Student’s unpaired t-test at any given lengthening contractions cycle and post lengthening contractions.

**Figure 5-figure supplement 1.**
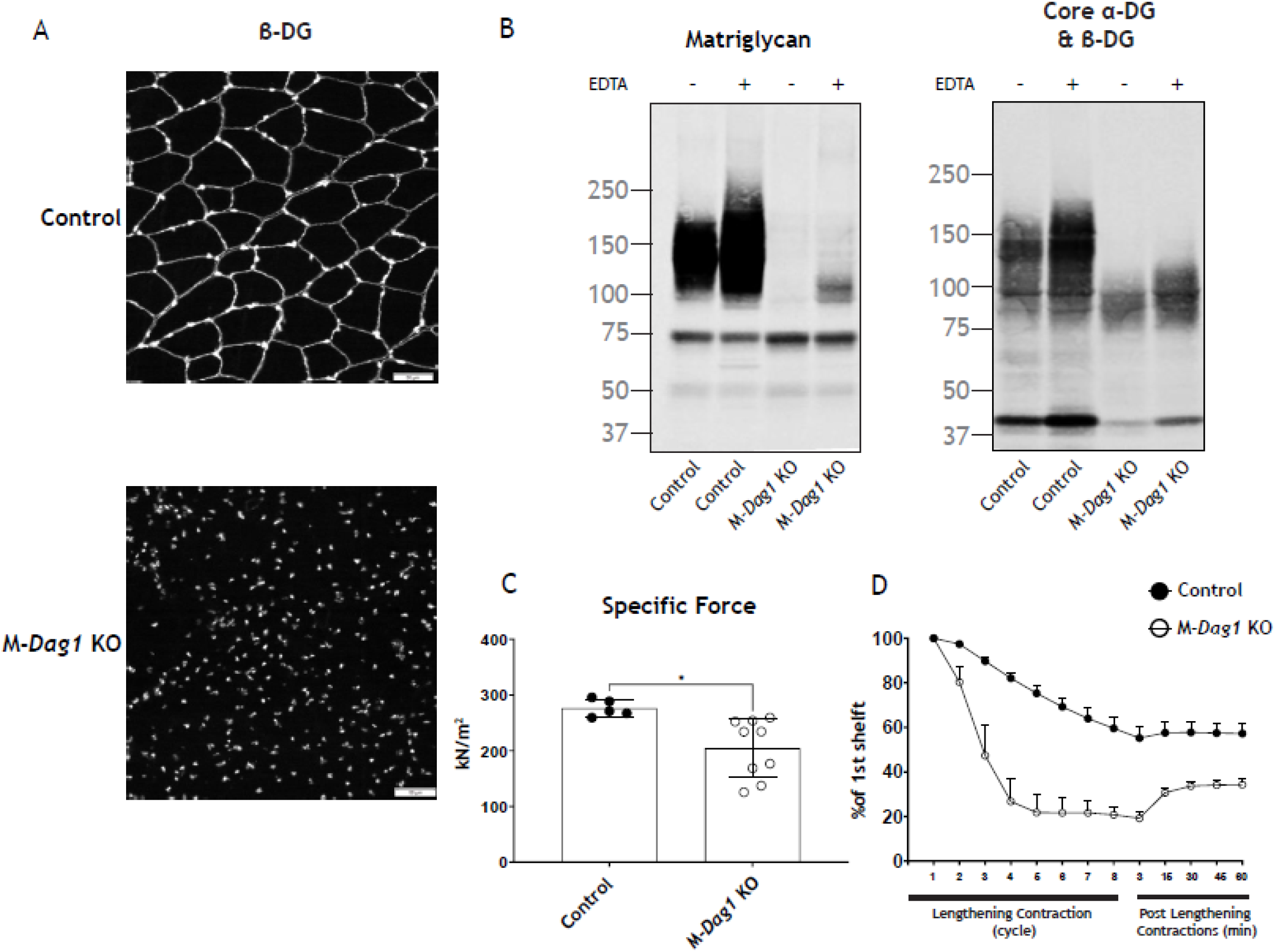
Characteristics of M-*Dag1* KO (*Pax7^cre^ Dag1^flox/flox^*) mice. **(A)** Immunofluorescence analyses of quadriceps muscles from a 12-week-old WT littermate (control) or M-*Dag1* KO mouse. Sections were stained to detect β-DG (AP83) and nuclei (DAPI). **(B)** Immunoblot analysis of skeletal muscle from control and M-*Dag1* KO mice. Glycoproteins were enriched from skeletal muscles (quadriceps) using WGA-agarose with (+) and without (-) 10 mM EDTA. Immunoblotting was performed to detect matriglycan (IIIH11) and core α-DG and β-DG (AF6868). **(C)** Specific force in EDL muscles of mice in indicated groups; p=0.0128, as determined by Student’s unpaired t-test. **(D)** Force deficit and force recovery after eccentric contractions in EDL muscles of 12- to 17-week-old male & female control (n=3) and M-*Dag1* KO (n=6) mice.

**Figure 6-figure supplement 1.**
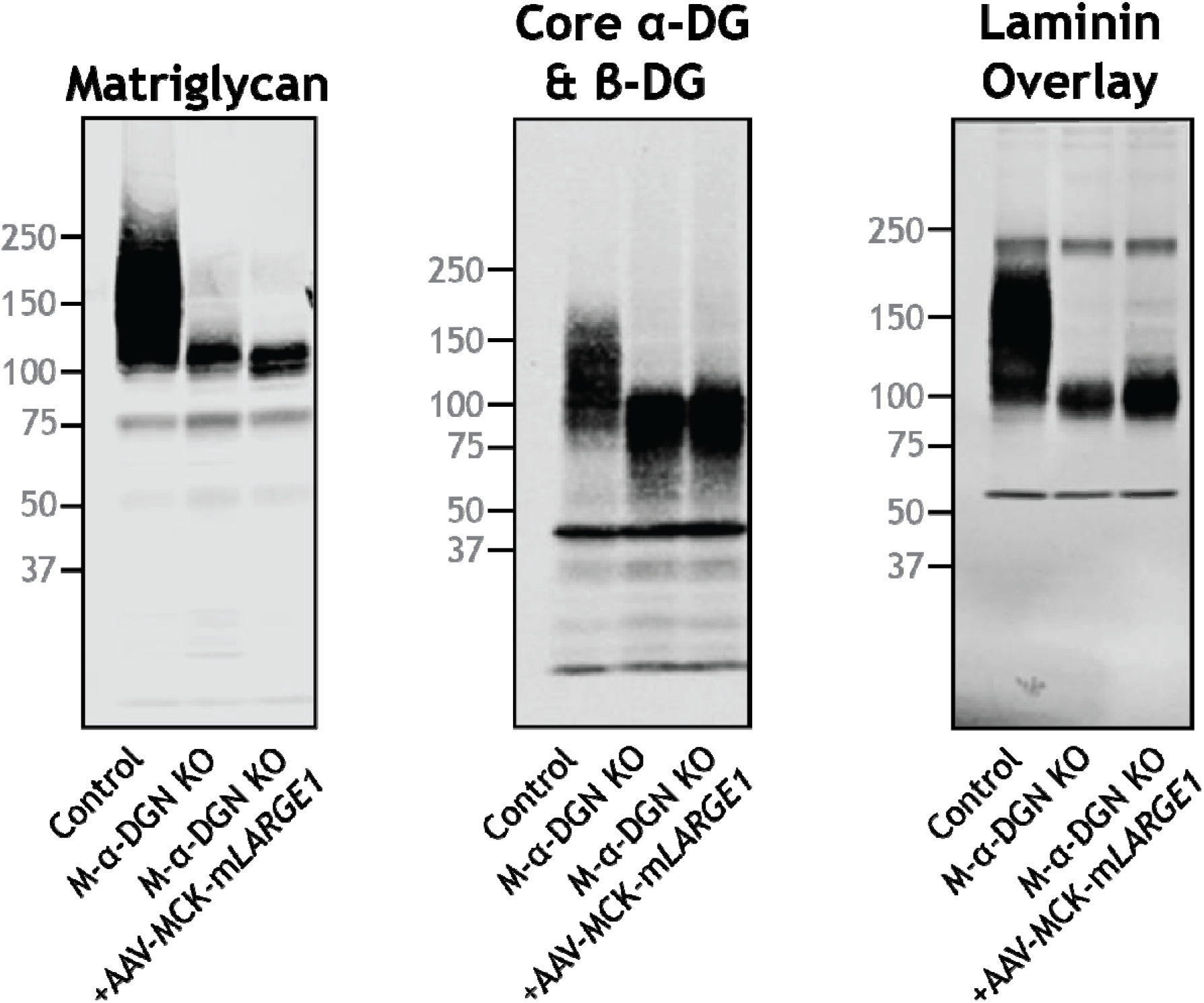
LARGE1 overexpression does not extend matriglycan on dystroglycan lacking α-DGN. *AAV-MCK-LARGE1* was injected into the retro-orbital sinus 10-to-24-week-old M-α-DGN KO mice. Quadriceps skeletal muscle was dissected 10 to 22 weeks after injection from control, M-α-DGN KO, and M-α-DGN KO+AAV-MCK-m*LARGE1* and used for immunoblotting analysis. Glycoproteins were enriched using WGA-agarose with 10 mM EDTA. Immunoblotting was performed to detect matriglycan (IIIH11), core α-DG and β-DG (AF6868), and laminin (overlay). Molecular weight standards in kilodaltons (kDa) are shown on the left.

**Figure 7-figure supplement 1.**
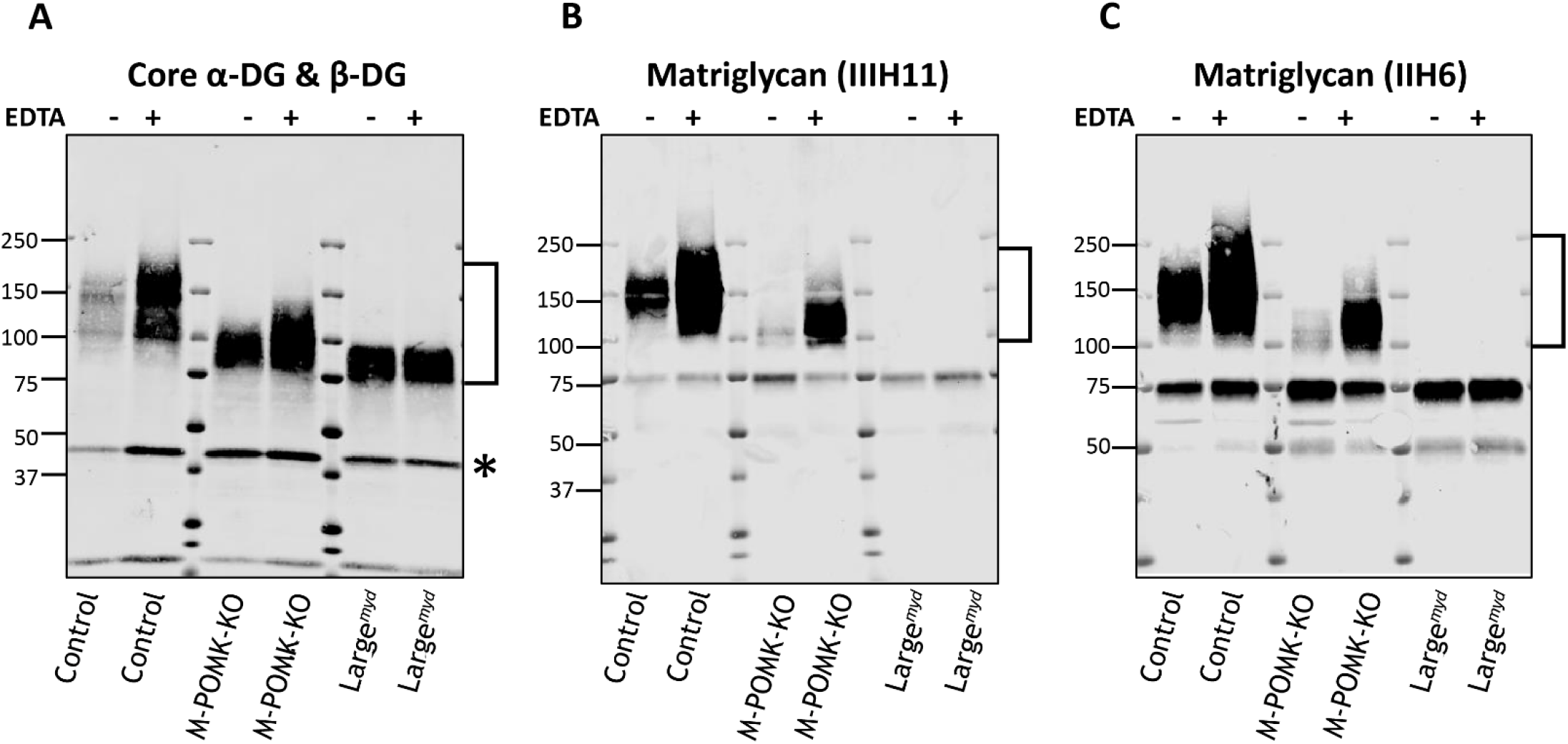
Effect of EDTA on solubilization of α-DG from skeletal muscle. Western blot analysis of DG in glycoprotein-enriched samples of control, M-POMK-KO, and *Large^myd^* skeletal muscle. Homogenates were prepared with (+) and without (-) 10mM EDTA (indicated on top) and enriched on WGA-agarose beads. Following washing, WGA-agarose beads were eluted with Laemmli sample buffer, and samples were loaded onto SDS-PAGE and blotted onto PVDF-FL membranes. Immunoblotting was performed to detect **(A)** core α-DG & β-DG (AF6868), **(B)** matriglycan (IIIH11) and **(C)** matriglycan (IIH6). α-DG is labeled with a bracket and varies in apparent molecular weight depending on glycosylation with matriglycan. β-DG is labeled with an asterisk.

